# Fanconi anemia-isogenic head and neck cancer cell line pairs - a basic and translational science resource

**DOI:** 10.1101/2022.09.11.507488

**Authors:** H. Tai Nguyen, Weiliang Tang, Andrew L.H. Webster, Jeffrey R. Whiteaker, Christopher M. Chandler, Ricardo Errazquin, Lucas B. Sullivan, Erica Jonlin, Elizabeth E. Hoskins, Eleanor Y. Chen, Madeline Fritzke, Amanda G. Paulovich, Susanne I. Wells, Khashayar Roohollahi, Josephine Dorsman, Ruud Brakenhoff, Ramon Garcia-Escudero, Agata Smogorzewska, Leslie Wakefield, Markus Grompe, Raymond J. Monnat

## Abstract

Fanconi anemia (FA) is a heritable malformation, bone marrow failure and cancer predisposition syndrome that confers an exceptionally high risk of developing carcinomas arising in squamous mucosal epithelia lining the mouth, proximal esophagus, vulva and anus. The origin of these cancers is not understood, and no effective way has been identified to prevent or delay their appearance. FA-associated carcinomas are also therapeutically challenging, as they may be multi-focal and stage-advanced at diagnosis making surgical control challenging. Moreover, individuals with FA have systemic DNA damage hypersensitivity and thus an elevated risk of toxicity when treated with standard-of-care therapies such as DNA cross-linking drugs and ionizing radiation.

We developed the Fanconi Anemia Cancer Cell Line Resource (FA-CCLR) in order to foster new research on the origins, treatment, and prevention of FA-associated cancers. The FA-CCLR consists of *FANC*-isogenic head and neck squamous cell carcinoma (HNSCC) cell line pairs from cancers arising in individuals with FA, or newly engineered from sporadic HNSCC cell lines. Molecular, cellular, and biochemical analyses were used to demonstrate the causal dependence of key FA-associated phenotypes on *FANC* genotype, expression and pathway activity. These *FANC*-isogenic cell line pairs are available to academic and non-profit investigators, with ordering information available at the ‘Fanconi Anemia Research Materials’ Resource and Repository at Oregon Health & Sciences University, Portland OR.

**Significance:** We have generated new isogenic cancer cell line models to investigate the origins, treatment and prevention of Fanconi anemia-associated squamous carcinomas that target the oral mucosa, proximal esophagus, and anogenital region.

## Introduction

Fanconi anemia (FA) is a heritable bone marrow failure, malformation and cancer predisposition syndrome that results from loss of function of any of the 23 functionally-linked Fanconi (*FANC*) genes (1,2). The initial cancer predisposition recognized in association with FA was leukemia, often in conjunction with bone marrow failure. This risk was substantially reduced, as was the risk of death due to bone marrow failure, by the development of highly effective bone marrow transplantation protocols. These protocols substantially changed the natural history of Fanconi anemia: there are now more adults living with FA than young individuals newly diagnosed with this syndrome.

An unanticipated consequence of the success in treating FA-associated bone marrow failure has been the emergence of another, unanticipated cancer risk: an extraordinarily high lifetime risk of developing squamous cell carcinomas (SCC) that arise in the mucosal epithelial lining the upper aerodigestive tract (oropharynx, neck and proximal esophagus), the anus and vulvar skin. Multiple oral cavity carcinomas are distressingly common (3), and an elevated risk of cutaneous squamous carcinomas has been documented together with a continuing, elevated risk of leukemia (3,4). However, SCC may be the first clinical presentation of FA at any age (1,5). Many other cancer types have been reported in individuals with FA, but only a small subset - liver and brain tumors, lymphomas, and embryonal tumors such as Wilms tumor and neuroblastoma - are observed at higher-than-expected frequencies versus population controls (3), and none with risk estimates approaching those of squamous mucosal carcinomas.

The origin of these FA-associated cancers is not well-understood. The loss of FA pathway function promotes constitutional genomic instability, and perturbs other cellular pathways that have the potential to elevate cancer risk (2,4,6). Smoking and excessive alcohol consumption appear to be less prominent drivers of HNSCC risk in individuals with FA than in the general population, but may contribute to risk as may prior bone marrow transplant conditioning (3). In contrast, human papillomavirus (HPV) infection, an important driver of SCC risk in the general population, has been detected in only a small subset of FA-associated anogenital SCC (7).

Effective therapies for FA-associated SCC have also proven elusive: many cancers are detected too late for surgical cure, and FA patients cannot tolerate effective doses of standard-of-care therapies including DNA cross-linking drugs and ionizing radiation (5). These features of cancer in FA - the often-late clinical recognition, synchronous or metasynchronous tumors and therapeutic intractability - have collectively made SCC the leading cause of premature morbidity and mortality in adults with FA.

In order to foster new work on the origins, treatment and prevention of FA-associated SCC, we developed a panel of *FANC*-isogenic SCC cancer cell line pairs from individuals with FA or engineered from sporadic, non-FA-associated SCC. All were derived from head and neck SCC (HNSCC) cell lines, the most common FA-associated SCC. The complementation groups FA-A, FA-C and FA-L encompass ∼75% of FA patients, and all three are represented among our FA patient-derived cell lines. Both the FA patient-derived and comparable newly engineered *FANCA*-mutant sporadic HNSCC cell lines were *FANC* transgene-complemented to generate isogenic cell line pairs. The generation and distribution of these *FANC*-isogenic cell line pairs to support research by academic and non-profit investigators have been generously supported by the Fanconi Anemia Research Fund, Eugene OR .

## Materials and Methods

### Cell line sources and culture

Twelve previously reported FA patient-derived or sporadic HNSCC cell lines were used to develop the FA-CCLR. All were given systematic names following International Cell Line Authentication Committee (ICLAC) guidelines (8) to allow unambiguous identification in conjunction with their uniquely defining DNA fingerprints. The human osteosarcoma-derived cell line U-2 OS was purchased from ATCC and used for methods development. Cell line identities were confirmed - or established - by the short tandem repeat (STR) genomic marker profiling of 9 standard human species-specific STR markers (CellCheck9 Plus, IDEXX BioAnalytics Westbrook, ME USA). Additional probes included with this panel were used to rule out interspecies contamination or *Mycoplasma ssp.* infection. Cell line use was considered ‘Not Human Subjects Research’ in light of NIH and University of Washington Institutional guidelines defining human subjects research. No patient-identifying information was associated with the resulting FA-CCLR cell lines.

FA-patient derived OHSU-SCC-974 and sporadic JHU-SCC-FaDu and UM-SCC-47 cell lines were grown in Eagle’s Minimum Essential Media (MEM) supplemented with 10% v/v FBS (Hyclone Laboratories, South Logan Utah) and 1% penicillin-streptomycin (Gibco #10378016). Sporadic cell line SFCI-SCC-9 was grown in a 1:1 mixture of Dulbecco-Modified Eagle’s Medium (D-MEM) and Nutrient Mixture F12 supplemented with 10% FBS, 0.5 mM sodium pyruvate, 400 ng/mL hydrocortisone, and 1% penicillin-streptomycin. All other cell lines were grown in D-MEM supplemented with 10% FBS, 1% non-essential amino acids (Gibco #11140050) and 1% penicillin-streptomycin unless otherwise noted. Cell growth was at 37°C in a 5% CO_2_/ambient (20%) oxygen water vapor-saturated incubator.

### Cell line genomic characterization

Prior reports and more recent, comprehensive genomic profiling data were used to identify cell line-specific single nucleotide variants (SNV), copy number variants (CNV) and cell line SNV/indel variant (9–13). Somatic SNV/indel and copy-number data for FA patient-derived cell lines OHSU-SCC-974, CCH-SCC-FA1 and CCH-SCC-FA2 were extracted from Webster et al. (12). In brief, somatic SNV/indel calls were made using CaVEMAN (14) and Pindel (15) using tumor cell line and patient matched normal control whole-genome sequencing (WGS) data.

Copy-number amplifications and deletions for OHSU-SCC-974, CCH-SCC-FA1 and CCH-SCC-FA2 were called using CNVkit (16) to compare tumor cell line and patient matched WGS data. SNV/indels and copy-number amplifications and deletions were called for FA patient-derived cell lines VU-SCC-1604, VU-SCC-1365 and VU-SCC1131 using, respectively, Mutect2 (17) and CNVkit to compare tumor cell line and patient matched fibroblast whole-exome sequencing (WES) data (13), with a focus on variant calling in significantly mutated genes considered to be HNSCC drivers when altered (9,18,19). Somatic mutation data for sporadic HNSCC cell lines JHU-SCC-FaDu, CAL-SCC-27, CAL-SCC-33, SFCI-SCC-09 and UM-SCC-01 were extracted from the Broad Institute Cancer Cell Line Encyclopedia (CCLE)(10), employing Mutect2 and Absolute (20) for SNV/indel and copy-number analyses. For all cell lines, the reporting threshold for copy-number amplifications and deletions was set at log2(CN)>0.5 or log2(CN)<-0.5 respectively.

### Generation and transgene complementation of FANC-deficient cell lines

*FANCA*-mutant sublines were developed in sporadic HNSCC by Cas9-mediated gene editing using dual guide RNA-targeted Cas9 deletion of a 5’ portion of *FANCA,* or single guide-targeted *FANCA* exon 4 cleavage to promote mutagenic end joining (21). Clonally-derived *FANCA*-mutant sublines were identified and characterized by colony deletion-specific PCR screening and/or Sanger sequencing prior to transgene complementation (21). *FANC* transgene complementations were performed by lenti- or retroviral transduction and by Cas9/Cas12a/CpfI-gRNA-targeted *FANC* transgene insertion into chromosome 4q Safe Harbor Site 231 (SHS231; (22)). Lentiviral low multiplicity-of-infection transductions utilized 3rd generation HIV-1-based *FANCA*, *FANCC*, *FANCF* or *FANCL* transgene vectors that had been CsCl gradient-purified and packaged according to Addgene-archived pLKO.1 protocols.

Retroviral transduction of the *FANCA* transgene vector S11FAIN was performed as previously described (21). Chromosomal SHS231 safe harbor site transgene cassettes were electroporated into *FANC*-deficient cells with expression plasmids encoding SHS231-specific gRNAs and Cas9 or Cas12a/Cpf1 nuclease. Transfected cells were grown for 3 days prior to adding hygromycin B (400μg/mL; Millipore Sigma, Catalog #400050) or puromycin (1µg/ml, Invivogen) for 10 - 14 days additional growth under selection prior to dilution cloning and screening to identify transgene-positive colonies and to verify both site-specific insertion and orientation for SHS231-safe harbor site-targeted transgenes.

### Cell proliferation

Cell population doubling times were quantified in different growth media to identify potential metabolic constraints as previously described (23). Cell cycle phase distributions were determined by 4′,6-diamidino-2-phenylindole (DAPI)/ethidium bromide staining of exponentially growing control or mitomycin-C-treated cultures. DNA content profiling was performed on a BD Biosciences LSR II Flow Cytometer using chicken erythrocytes as a DNA standard, followed by cell cycle phase quantification using FlowJo™ software (FlowJo, Ashland, OR USA) as previously described (24).

### Drug and small molecule response profiling

Drug and small molecule dose- and genotype-dependent sensitivity was determined for the chemotherapeutic drugs *cis*-Pt, carboplatin, and oxaliplatin; for formaldehyde; for the kinase inhibitors gefitinib and afatinib; inhibitors of the ATR kinase (M6620, VX-970/VE822) and WEE1 (AZD-1775); and for metformin and rapamycin. Two representative FA patient-derived and two sporadic isogenic cell line pairs were used for most analyses. In brief, triplicate wells containing 2,500 cells/well were seeded in 96-well plates in 150 µL of complete growth medium, followed by the addition of two- or three-fold serial dilutions of drugs or small molecules in 50 μl of medium, followed by 4 days of growth in the presence of drug/small molecule. Cell survival was then quantified versus a DMSO/vehicle control by WST-1 (Takara Bio, Catalog #MK400) or AlamarBlue (ThermoFisher, Catalog #DAL1025) assay. Data were normalized versus control (mock-treated/vehicle-only treated) wells, and plotted using GraphPad PRISM (GraphPad Software, San Diego, CA) to determine dose-dependent growth suppression and to estimate drug/small molecule IC_50_ values.

### Protein expression analyses

Western blot analyses of FANC protein expression were performed using total cell protein extracts (∼40 μg from ∼1-2.5 × 10^6^ cells) prepared in RIPA Lysis Buffer (Boster Biological Technology, Pleasanton CA, Catalog #AR0105) containing protease inhibitors as previously described (25). Proteins were size-fractionated on precast 10 cm 4-12% Bis-Tris or Tris-acetate gels (Thermo Fisher Catalog #NP0336BOX), then electroblotted onto nitrocellulose membranes (Thermo Fisher Catalog #88025). Membranes were blocked in Tris-buffered saline solution containing 5% non-fat dry milk for 1 hr at room temperature (or overnight at 4°C), then incubated with FANC protein-specific rabbit polyclonal or mouse monoclonal primary antibodies detected using a horseradish peroxidase-conjugated anti-rabbit IgG goat or anti-mouse antibody. All transgenes apart from *FANCA* could also be detected by blotting for an in-frame, N-terminal HA epitope tag using an anti-HA epitope tag-specific mouse monoclonal antibody. All antibodies, suppliers and protocols are detailed in Extended Methods.

Peptide immunoenrichment coupled with targeted multiple reaction monitoring mass spectrometry (immuno-MRM) was also used to detect and quantify the expression and modification status of 66 DNA damage response (DDR) proteins including 10 FANC proteins. In brief, triplicate 10 cm dishes were seeded with 2 - 5 × 10^6^ cells prior to the addition of 0.2 µM mitomycin C for 24 hrs. Whole cell lysates were prepared and processed in a blinded fashion as previously described, using two antibody panels that targeted 126 peptides and 59 post-translational modification sites (26–28). MRM-MS data were analyzed using Skyline (29), with manual review of peak integrations to confirm and align peptide transitions of endogenous and stable mass isotope-labeled peptide standards. Means and standard deviations of peak area ratios were calculated from three complete biological replicates of cell line ± treatment, then combined and reported in a single list of analytes.

### In vitro three-dimensional ‘tumoroid’ cultures

Three-dimensional (3D) ‘tumoroid’ cultures were grown from *FANC*-isogenic HNSCC JHU-SCC-FaDu and VU-SCC-1131 cell line pairs by modifying a previously reported protocol (30). In brief, single cell suspensions were plated in Matrigel (Corning Life Sciences, Durham, NC USA) to form 40µl ‘domes’ each containing ∼1000 cells that were grown for 7 days submerged in media before being gently dissociated for additional growth in suspension. MMC responsiveness was assessed by growing Day 40 tumoroids for 3 additional days in medium supplemented with 10, 50 or 100 nM MMC. Tumoroid samples were then fixed and paraffin embedded for H&E staining, and immunostaining to detect cytokeratin 5 (CK5), the p40 fragment of p63 and the functional state markers Ki67, Ser139-H2AX phosphorylation and cleaved caspase 3.

### Statistical analyses

Cell proliferation and dose-dependent survival analysis are displayed with mean ± the standard error of the mean (SEM) values. The statistical significance of differences in cell proliferation rate as a function of genotype was determined by one-way ANOVA. The data shown in the cell cycle analysis represent the phase fractions extracted from a minimum of 30,000 gated and flow-analyzed single cells.

### Availability of materials and data

The HNSCC isogenic cell line pairs generated as part of this project were expanded, re-authenticated and verified to be *Mycoplasma-*free prior to distribution. Information including isogenic cell line pair-specific data sheets, and availability for academic and non-profit investigators can be found at the Fanconi Anemia Research Materials (FARM) Resource website (https://apps.ohsu.edu/research/fanconi-anemia/cell_line/). The FARM Resource and this project were generously supported by the Fanconi Anemia Research Fund, Eugene OR. Additional data generated or used as part of this project that is not in the public domain can be requested from the corresponding author.

## Results

A systematic workflow was used to generate isogenic cell line pairs from FA patient-derived and newly generated *FANCA-*mutant, sporadic HNSCC cell lines (Figure 1). The FA patient-derived cell lines were from HNSCC arising in FA patients from complementation groups FA-A, FA-C and FA-L, with causative mutations in, respectively, the *FANCA*, *FANCC* and *FANCL* genes. Sporadic HNSCC *FANC*-isogenic cell line pairs were generated by gene editing to inactivate *FANCA,* followed by *FANCA* transgene complementation (Table 1). All cell line identities were confirmed by short tandem repeat DNA ‘fingerprinting’, and verified to be free of cross-species contamination or *Mycoplasma* infection (Table S1).

**Figure 1:**
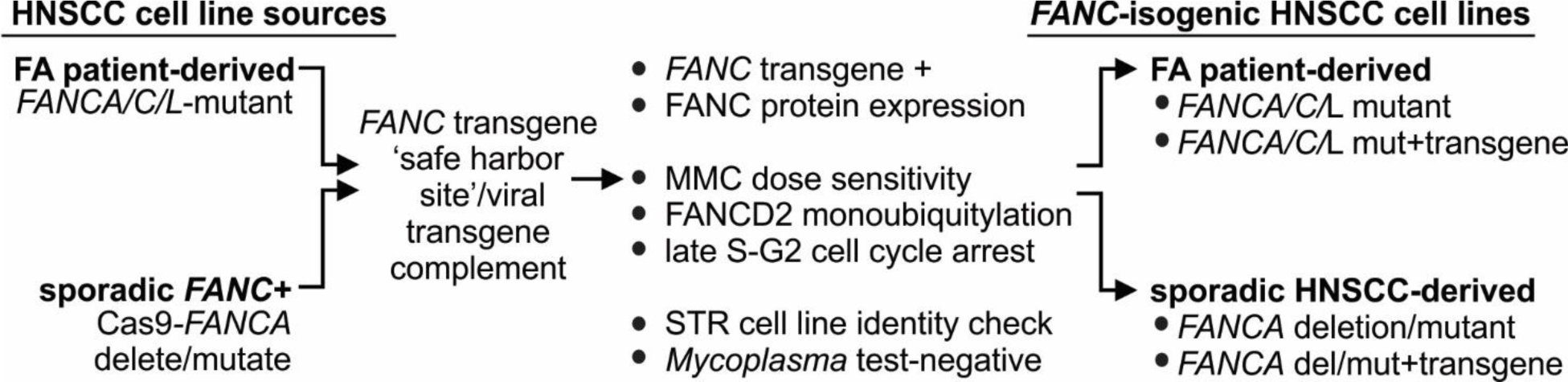
Workflow to generate FANC-isogenic cell line pairs from FA patient-derived and sporadic head and neck cancer cell lines.

**Table 1.**
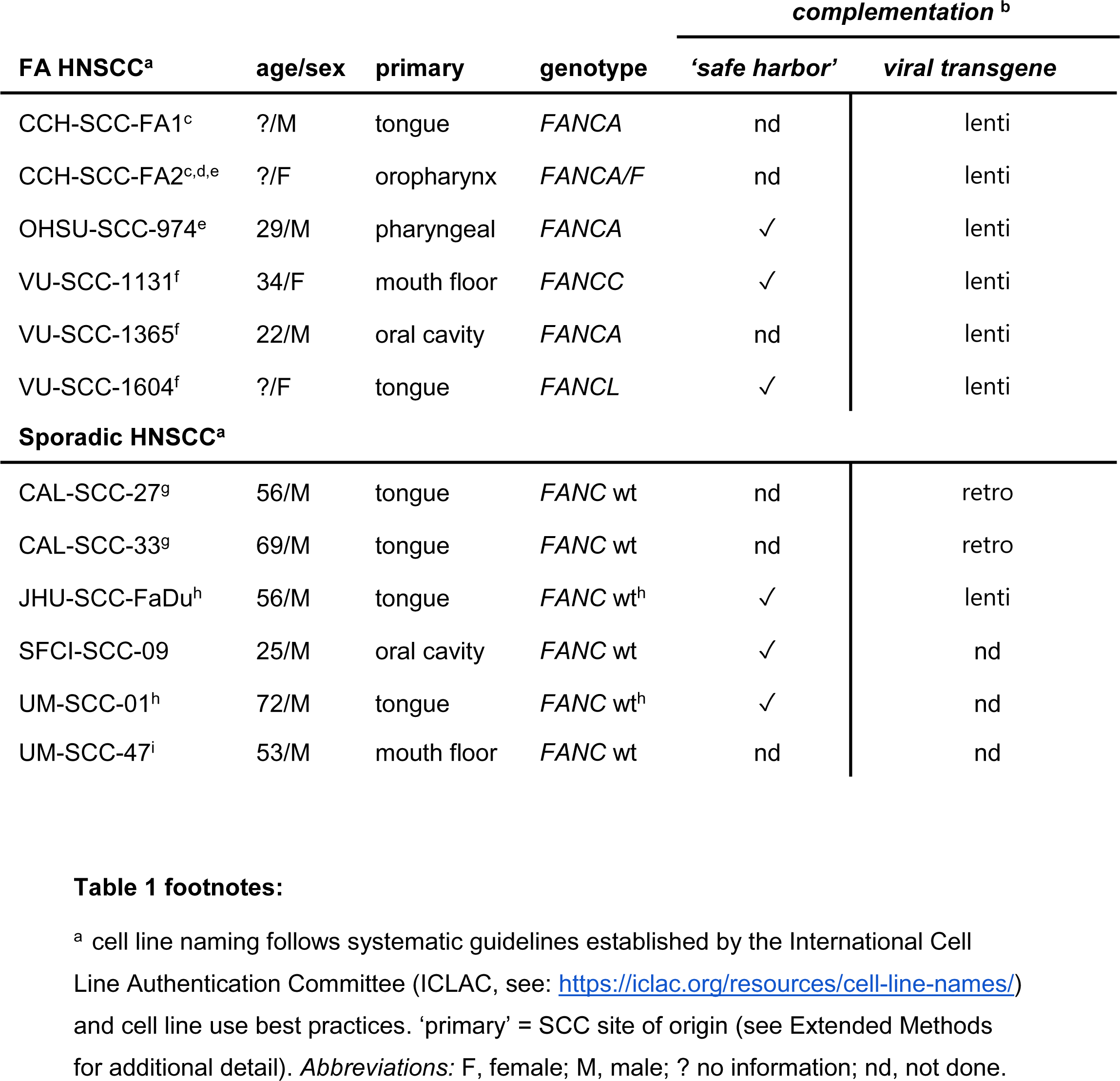
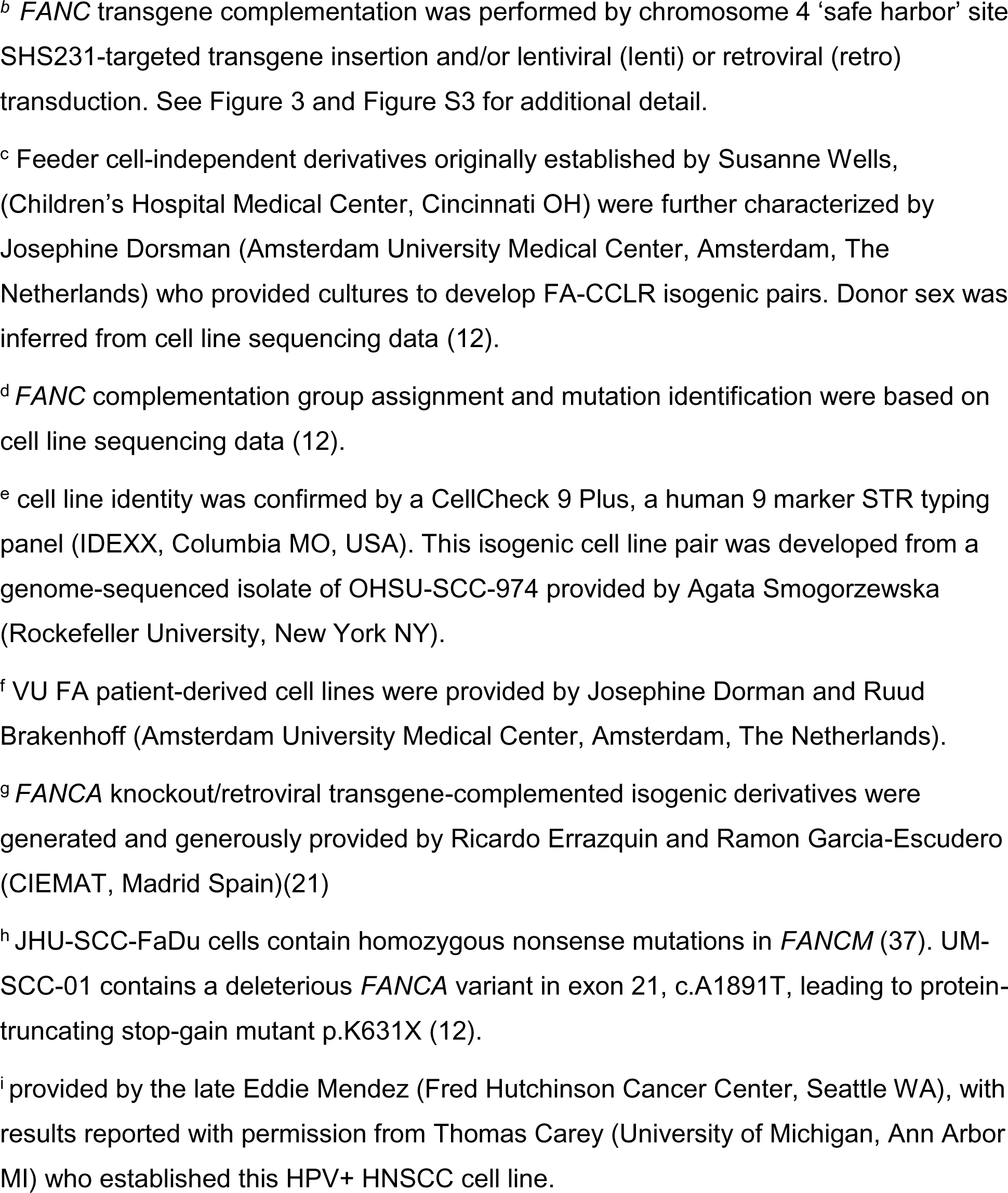
Fanconi anemia patient and sporadic head and neck cancer cell lines.

| FA HNSCC<sup>a</sup> | age/sex | primary | genotype | *complementation*<sup>b</sup> |  |
| --- | --- | --- | --- | --- | --- |
|  |  |  |  | 'safe harbor' | viral transgene |
| CCH-SCC-FA1<sup>c</sup> | ?/M | tongue | *FANCA* | nd | lenti |
| CCH-SCC-FA2<sup>c,d,e</sup> | ?/F | oropharynx | *FANCA/F* | nd | lenti |
| OHSU-SCC-974<sup>e</sup> | 29/M | pharyngeal | *FANCA* | ✓ | lenti |
| VU-SCC-1131<sup>f</sup> | 34/F | mouth floor | *FANCC* | ✓ | lenti |
| VU-SCC-1365<sup>f</sup> | 22/M | oral cavity | *FANCA* | nd | lenti |
| VU-SCC-1604<sup>f</sup> | ?/F | tongue | *FANCL* | ✓ | lenti |
| **Sporadic HNSCC<sup>a</sup>** |  |  |  |  |  |
| CAL-SCC-27<sup>g</sup> | 56/M | tongue | *FANC* wt | nd | retro |
| CAL-SCC-33<sup>g</sup> | 69/M | tongue | *FANC* wt | nd | retro |
| JHU-SCC-FaDu<sup>h</sup> | 56/M | tongue | *FANC* wt<sup>h</sup> | ✓ | lenti |
| SFCI-SCC-09 | 25/M | oral cavity | *FANC* wt | ✓ | nd |
| UM-SCC-01<sup>h</sup> | 72/M | tongue | *FANC* wt<sup>h</sup> | ✓ | nd |
| UM-SCC-47<sup>i</sup> | 53/M | mouth floor | *FANC* wt | nd | nd |

<sup>a</sup> cell line naming follows systematic guidelines established by the International Cell Line Authentication Committee (ICLAC, see: https://iclac.org/resources/cell-line-names/) and cell line use best practices. 'primary' = SCC site of origin (see Extended Methods for additional detail). *Abbreviations*: F, female; M, male; ? no information; nd, not done.
<sup>b</sup> *FANC* transgene complementation was performed by chromosome 4 'safe harbor' site SHS231-targeted transgene insertion and/or lentiviral (lenti) or retroviral (retro) transduction. See Figure 3 and Figure S3 for additional detail.
<sup>c</sup> Feeder cell-independent derivatives originally established by Susanne Wells, (Children's Hospital Medical Center, Cincinnati OH) were further characterized by Josephine Dorsman (Amsterdam University Medical Center, Amsterdam, The Netherlands) who provided cultures to develop FA-CCLR isogenic pairs. Donor sex was inferred from cell line sequencing data (12).
<sup>d</sup> *FANC* complementation group assignment and mutation identification were based on cell line sequencing data (12).
<sup>e</sup> cell line identity was confirmed by a CellCheck 9 Plus, a human 9 marker STR typing panel (IDEXX, Columbia MO, USA). This isogenic cell line pair was developed from a genome-sequenced isolate of OHSU-SCC-974 provided by Agata Smogorzewska (Rockefeller University, New York NY).
<sup>f</sup> VU FA patient-derived cell lines were provided by Josephine Dorman and Ruud Brakenhoff (Amsterdam University Medical Center, Amsterdam, The Netherlands).
<sup>g</sup> *FANCA* knockout/retroviral transgene-complemented isogenic derivatives were generated and generously provided by Ricardo Errazquin and Ramon Garcia-Escudero (CIEMAT, Madrid Spain)(21)
<sup>h</sup> JHU-SCC-FaDu cells contain homozygous nonsense mutations in *FANCM* (37). UM-SCC-01 contains a deleterious *FANCA* variant in exon 21, c.A1891T, leading to protein-truncating stop-gain mutant p.K631X (12).
<sup>i</sup> provided by the late Eddie Mendez (Fred Hutchinson Cancer Center, Seattle WA), with results reported with permission from Thomas Carey (University of Michigan, Ann Arbor MI) who established this HPV+ HNSCC cell line.

FA-CCLR cell lines displayed both HNSCC- as well as FA-associated genomic alterations (9,12,13,31–33). Figure 2 summarizes cell line-specific single nucleotide (SNV) and copy number (CNV) variant data for cell lines where we had full data access and compares these lines with HPV-negative HNSCC samples included in TCGA data (n = 426; (19)). We had limited data access for the three VU FA patient-derived cell lines, and thus focused on calling variants in a curated panel of genes frequently or significantly mutated in and considered potential drives of HNSCC (18,19). *TP53* variants were present in nearly all cell lines, together with different combinations of deleterious variants in HNSCC significantly-mutated and/or putative driver genes (Figures 2 and S1). None of the FA patient-derived cell lines contained identifiable HPV-derived DNA sequences, and consistent with previous reports (12) the vast majority of gene alterations in FA patient-derived HNSCC cell lines were gene deletion/ amplification events as opposed to basepair-level alterations.

**Figure 2.**
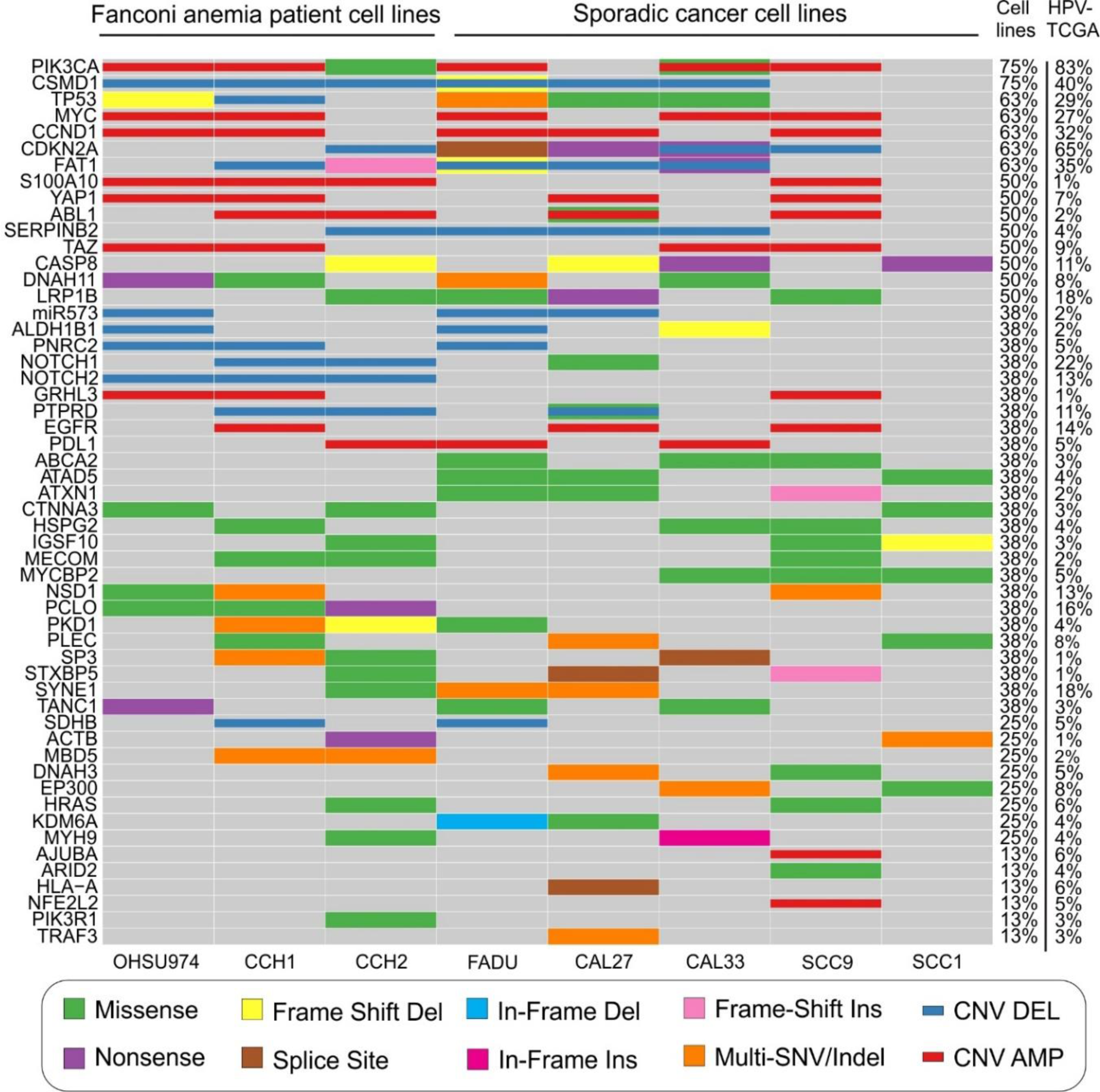
– Mutational landscape of Fanconi anemia patient & sporadic HNSCC cell lines. Oncoplot of eight head and neck cancer cell lines, including Fanconi anemia patient-derived SCC lines OHSU-SCC-974, CCH-SSC-FA1 and CCH-SCC-FA2, together with five sporadic HNSCC cell lines (JHU-SCC-FaDu, CAL-SCC-27, CAL-SCC-33, SFCI-SCC-09 and UM-SCC1-01. Displayed along the left margin is a curated panel of oncogenes and tumor suppressor genes frequently mutated in SCC. The Figure key at bottom indicates the type of mutation identified, while the two columns along the right margin indicate the frequency with which each gene was altered in SCC cell lines versus a comparison dataset of HPV-negative HNSCC included in TCGA. The reporting thresholds for copy-number amplifications and deletions are log2(CN)>0.5 or log2(CN)<-0.5 respectively. Copy-number data was unavailable or SCC1. See Table 1 for additional cell line information.

### Generation of FANC-isogenic HNSCC cell line pairs

Five independent sporadic HNSCC cell lines were used to generate *FANCA*-mutant and complemented isogenic cell line pairs for comparison with FA patient-derived isogenic cell lines. *FANCA-*mutant sporadic sublines were generated by Cas9 dual guide RNA-targeted deletion of *FANCA* exon 2 and a portion of exon 3, or by Cas9 single guide RNA-targeted cleavage of *FANCA* exon 4 to promote mutagenic end-joining. Resulting *FANCA* mutations were verified by a combination of PCR deletion or mismatch cleavage screening, DNA sequencing and Western blot analysis, to confirm the presence of allelic mutations and the loss of FANCA protein expression (Figure 3A and 3B)(21). We were able to generate only single allele *FANCA* deletions in the HPV+ sporadic HNSCC cell line UM-SCC-47. This result may reflect a potential synthetic-sick or -lethal interaction between HPV and the loss of FA function in this cell line, and is paralleled by the absence of HPV+ samples among 55 genomically characterized FA-associated HNSCC (12).

**Figure 3.**
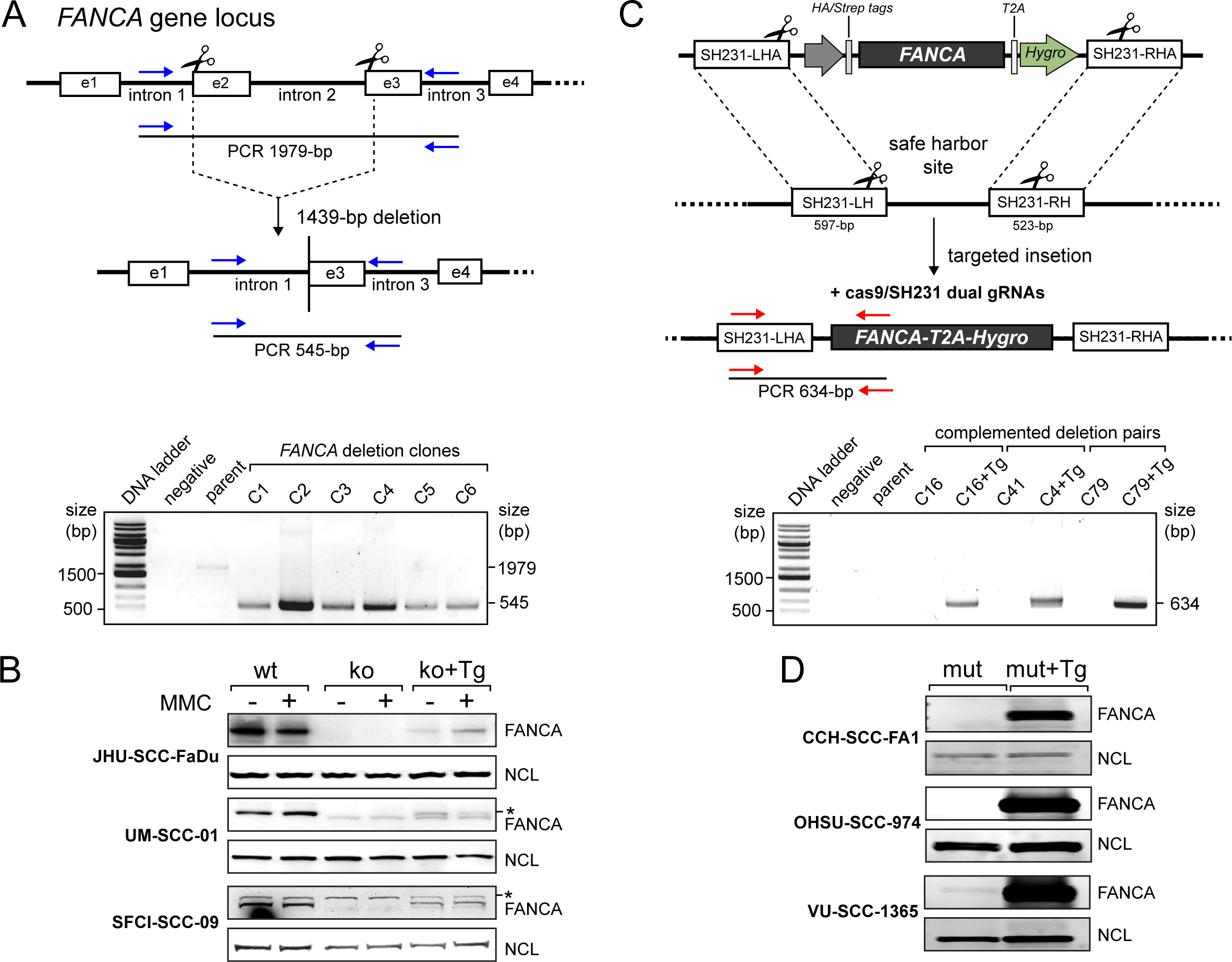
Generation and transgene complementation of *FANC*-deficient HNSCC cell lines. **A.** FANCA inactivation by dual guide RNA-targeted biallelic *FANCA* 5’ gene deletion (upper panel) with PCR detection of intact and deleted alleles (lower panel). **B.** FANCA protein expression in sporadic HNSCC cell lines JHU-SCC-FaDu, UM-SCC-01, and SFCI-SCC-09 prior to and after transgene insertion, and in mutant parent and *FANCA* transgene-complemented cells prior to and after MMC treatment. **C.** *FANCA* transgene insertion into chromosome 4q safe harbor site SHS231 by homology-mediated break repair (upper panel), with transgene insertion and orientation were confirmed by site- and orientation-specific PCR assay (lower panel). **D.** FANCA protein expression in FA patient-derived and *FANCA* transgene-complemented cell lines CCH-SCC-FA1, OHSU-SCC-974 and VU-SCC-1365. *Abbreviations:* LHA, left homology arm; RHA, right homology arm; LH, left homology; RH, right homology; wt = wild-type; mut = mutant; ko = gene knockout line; Tg = transgene; NCL = nucleolin, a loading control. * = non-specific band variably detected by FANCA antiserum.

*FANC*-complemented derivatives of FA patient-derived and newly-generated *FANCA*-mutant sporadic cell lines were generated by lenti- or retroviral transduction or by chromosome 4 safe harbor site SHS231 transgene insertion (Figure 3C). Viral complementation led to consistently higher transgene protein expression than did safe harbor site targeting, and was required to fully restore MMC resistance in FA patient-derived - but not newly generated sporadic, *FANCA*-mutant - HNSCC cell lines. Mutant/deleted and transgene-complemented cell line clones were comparable to parental cell lines or transgene-complemented cell pools, indicating their functional equivalence as assessed by MMC dose-survivals. Figure S2 shows one example of these analyses, in which MMC dose-dependent survival of patient-derived *FANCA-* mutant OHSU-SCC-974 cells was compared with the patent cell line, 11 independent transgene-complemented clonal derivatives and a pool of lentiviral transgene-complemented cells. These and additional results led us to distribute transgene-complemented pools of *FANC*-deficient cells with clonal complexities of ≥200 as a versatile starting point for most anticipated uses and end users. We were unable to generate a convincingly complemented isogenic cell line pair that restored MMC resistance for the FA patient-derived cell line CCH-SCC-FA2 that contains biallelic *FANCA* mutations and apparent *FANCF* gene silencing (Table 1). None of the complementation strategies we attempted – sequential single gene or dual gene transfer, safe harbor site insertion or viral transduction - fully restored MMC resistance for unknown reasons.

### Characterization of FANC-isogenic HNSCC cell line derivatives

Key *FANC* gene- and pathway activity-dependent phenotypes were characterized in order to establish the causal relationship linking cell line genotype and FANC protein expression to cellular phenotype. FA-dependent phenotypes we investigated were cell proliferation rate; *FANC* transgene expression; MMC dose-dependent cell killing and the induction of FANCD2 monoubiquitylation detected by Western blot and/or mass spectrometry; and MMC induction of a late S/G_2_. cell cycle arrest. A proliferative advantage of FANC-proficient cells has long been recognized in specific cell lineages both *in vivo* and *in vitro,* and this observation has provided the strong rationale for hematopoietic stem cell FA gene therapy to prevent bone marrow failure (34). We observed clear cell line- and growth medium-specific differences among *FANC*-isogenic cell line pairs, though these differences did not clearly segregate with *FANC* genotype. Comparable results have been reported by Errazquin et al. is FANCA-mutant CAL-SCC-27 and CAL-SCC-33 cells (21). Figure 4A provides a representative example of the proliferation of two *FANC*-isogenic patient-derived cell line pairs in different growth media at 37°C in ambient (20%) oxygen. Proliferation rates varied by over 2-fold, ranging from 0.47 - 1.2 doublings/day, with FA patient-derived cell line VU-SCC-1131 displaying the fastest and FA patient-derived cell line VU-SCC-1365 the slowest doubling rates. We did not observe genotype-specific proliferation rate differences between cells grown in ambient (20%) versus low (5%) oxygen (additional results not shown).

**Figure 4.**
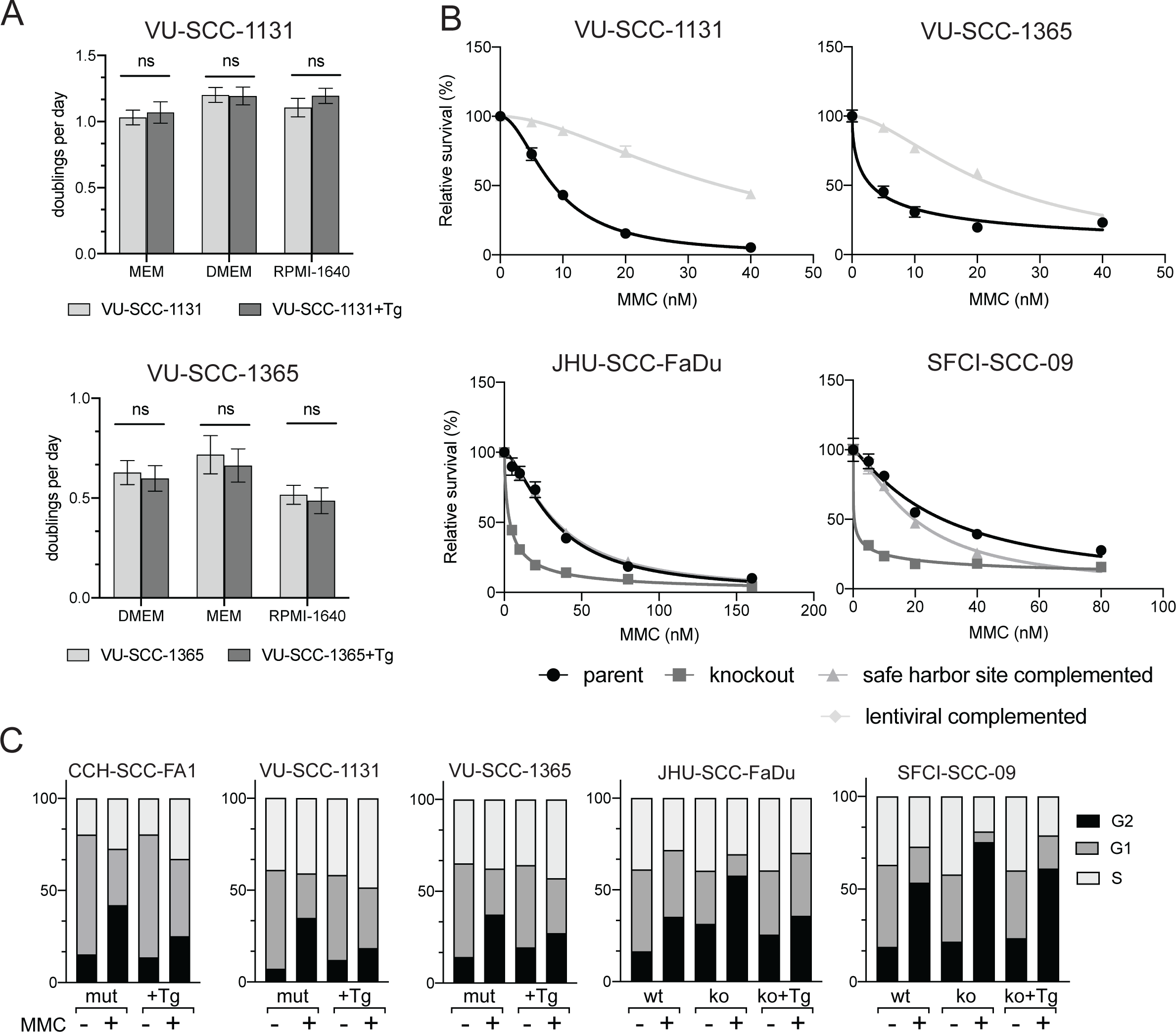
FA-dependent phenotypes in *FANC*-isogenic HNSCC cell lines. **A.** Comparative proliferation rates of *FANCC*-isogenic FA patient-derived (VU-SCC-1131) and *FANCA*-isogenic FA patient-derived (VU-SCC-1365) HNSCC cell line pairs in three different growth media. **B.** Mitomycin-C dose-dependent survival of isogenic FA patient-derived (VU-SCC-1131 and VU-SCC-1365) and sporadic (JHU-SCC-FaDu and SFCI-SCC-09) HNSCC cell line pairs. **C.** Cell cycle phase distributions for isogenic FA patient-derived CCH-SCC-FA1, VU-SCC-1131, and VU-SCC-1365 and sporadic JHU-SCC-FaDu and SFCI-SCC-09 HNSCC cells prior to MMC treatment, and MMC treated after *FANCA* gene deletion/transgene complementation. N = 30,000 events; Data represents mean ± SEM.

FA-deficient cells are exquisitely sensitive to MMC-induced cell killing and to an MMC-induced late S/G_2_ cell cycle arrest. These cellular phenotypes are so consistently robust that they remain cornerstones in FA diagnosis and mechanistic analyses. In contrast to cell proliferation, *FANC* genotype modified MMC sensitivity in a highly reproducible way in all isogenic cell line pairs, though again with substantially different cell line-specific differences (Table S2). FA patient-derived cell lines were predictably MMC-sensitized, with MMC IC_50_ values of 2.8 - 13.4 nM (mean 6.5 nM), ∼ 4-fold *lower* than FA-proficient sporadic HNSCC cell lines (range 4.4 - 36.8 nM, mean 22.9 nM). *FANC* transgene complementation of FA patient-derived cell lines conferred MMC resistance, with 3-to-10-fold *higher* MMC IC_50_ values. In contrast, *FANCA* deletion in sporadic HNSCC cell lines sensitized cells to MMC, with 14-to-44-fold *lower* MMC IC_50_ values (Table S2 and Table 1 of (21)).

MMC treatment of FA-deficient cells also provokes a consistent late S/G_2_ cell cycle arrest that has been useful for FA clinical diagnosis and mechanistic analyses (35). FA patient-derived cell lines displayed strong MMC dose-dependent increases in late S/G_2_ cell cycle fractions after treatment with 10 nM or 40 nM MMC. This arrest phenotype, most easily quantified by flow cytometry as an elevated G_2_/M fraction, revealed G_2_/M fraction that ranged from 16.4% (range 13.1% - 25.2%) in untreated cells, up to 41.3% (range 23.6 - 65.7%) in cells treated with 40 nM MMC. *FANC* transgene complementation had little effect on G_2_/M fractions in untreated cells, though substantially reduced G_2_/M fractions in MMC-treated cells. Comparable results have been reported by Errazquin et al. who used diepoxybutane (DEB), another DNA-damaging agent that selectively kills FA-deficient cells, to treat *FANCA*-mutant derivatives of CAL-SCC-27 and CAL-SCC-33 cells (21).

MMC treatment also induces the monoubiquitylation of FANCD2, a key regulatory post-translational modification and biomarker of FA pathway activity (2). This activity, mediated by the FA core complex protein and E3 ubiquitin ligase FANCL, is lost in ∼ 90% of FA patients (36). In order to further investigate this biochemical phenotype, and the proteomic response to the loss and restitution of FA pathway functional integrity, we employed peptide immunoenrichment coupled with targeted multiple reaction monitoring mass spectrometry (immuno-MRM). This approach allowed us to simultaneously detect and quantify FANCD2 expression and monoubiquitylation status, together with the abundance and post-translational modification status of peptides located in 65 additional DNA damage response proteins including 9 other FANC proteins. The outline of these experiments is shown in Figure 5A (26–28), in which we were able to quantify 98 of 129 potential target peptides using experimentally-defined lower limits for quantification (LOQ)(Figure 5B and 5C, Table S3).

**Figure 5.**
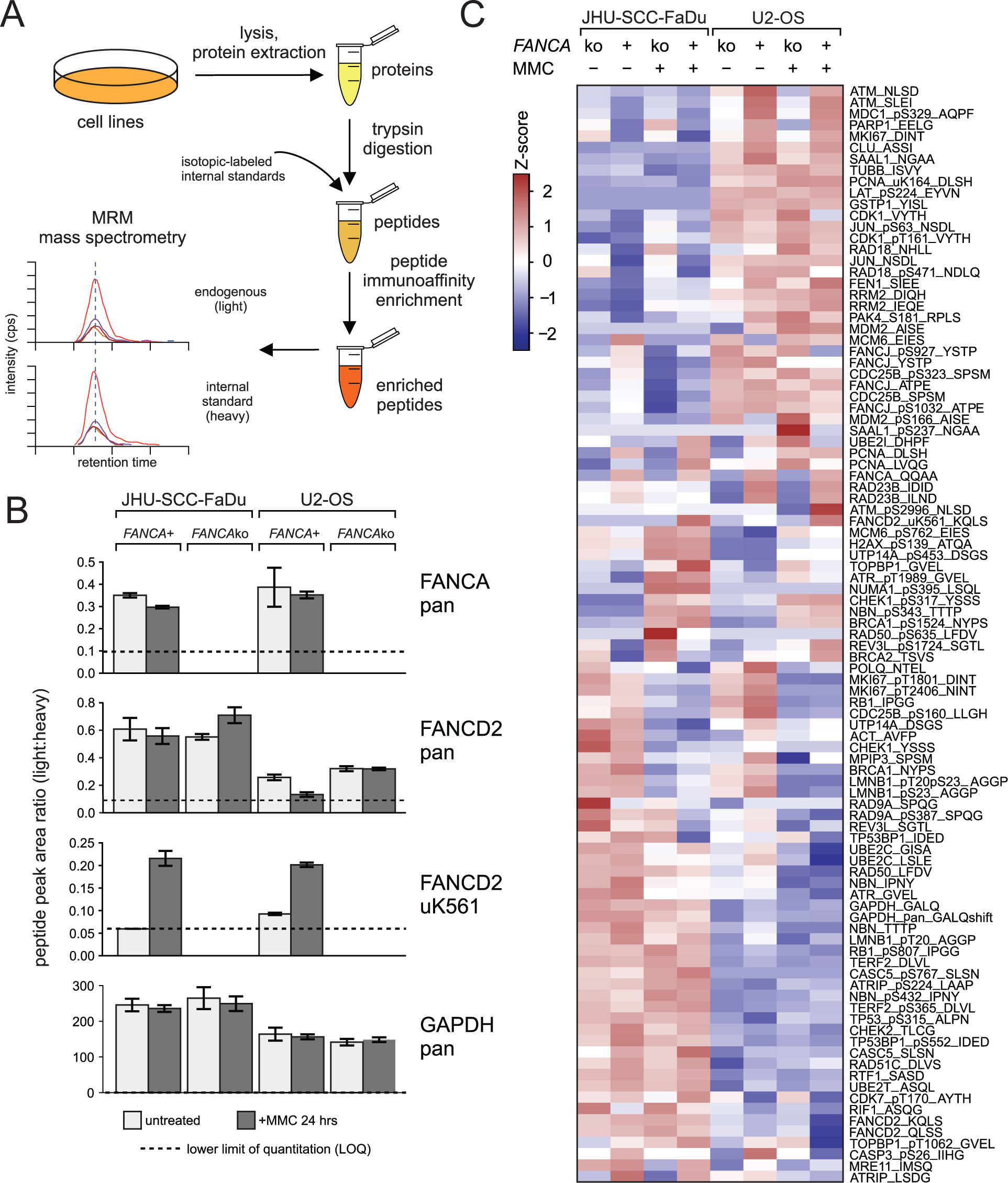
Immuno-MRM analysis of FANCA protein expression and FANCD2 ubiquitylation in response to MMC-mediated DNA damage. **A**, Experimental outline for sample handling with inclusion of mass labeled internal standard peptides for MRM mass spectrometry. **B,** Bar plots of peak area ratios (light:heavy peptide) for FANCA, FANCD2 and FANCD2 ubiquitination (uK561). GAPDH is a protein loading control. Error bars show the standard deviation of triplicate analysis. The dashed line indicates the experimentally-determined lower limit of quantification (LOQ). **C,** Heatmap showing unsupervised clustering of 100 analyte peptides detected above LOQ in JHU-SCC-FaDu and U2-OS cells after 24 h of MMC. Peak area ratios (light:heavy) were normalized for each peptide analyte. A subset of peptides near the center of the heatmap show coordinate increases or decreases in abundance in response to MMC treatment.

FANCA protein was readily detected in parental JHU-SCC-FaDu and control U-2 OS cells, but not in *FANCA*-mutant isogenic cell line derivatives. FANCD2 protein expression was unchanged in *FANCA-*proficient or mutant cells (Figure 5B), whereas FANCD2 ubiquitylation, detected as a ubK561-modified peptide, was induced in both parental cell lines but lost in both *FANCA-*mutant cell backgrounds (Figure 5B). Our ability to detect FANCD2 ubiquitylation by immuno-MRM, though not consistently by Western blot, in nominally *FANCA-*proficient JHU-SCC-FaDu cells, may reflect in part homozygous *FANCM* nonsense mutations in these cells (37) with the potential to attenuate the MMC-induced DNA damage response without conferring a clear FA-deficient cellular phenotype.

Other functionally important DNA damage-dependent post-translational modifications (e.g., ATRpT1989, H2AXpS139/γH2AX and CHEK1pS317) could be detected in the presence of FANCA loss, together with additional cell line-specific protein expression and/or modification differences that drove prominent sample clusters of MMC treatment-responsive peptides in both cell line backgrounds (Figure 5C, Table S3). These results demonstrate the ability of immuno-MRM to quantify expression and the DNA damage-dependent modification of many proteins in parallel in response to DNA damage or other perturbations.

### Isogenic cell line responses to additional drugs and small molecules

We determined the ability of nine additional drugs or small molecules in addition to MMC to suppress cell proliferation as a function of *FANC* genotype and dose. These additional agents were chosen on the basis of mechanism of action, and/or prior reports of *FANC* genotype-dependent activity. The loss of FA pathway activity did not consistently sensitize FA patient-derived or sporadic, *FANCA*-mutant HNSCC lines to formaldehyde dose-dependent cell killing over a 4 day/continuous exposure time course (Figure S4). This may reflect cell lineage-specific or other differences that collectively determine formaldehyde toxicity (38,39). In contrast, loss of FA pathway function sensitized cells regardless of origin to platinum-based chemotherapeutic drugs with *cis*-Pt displaying the strongest - and oxaliplatin the least – dose-dependent reductions in survival. Comparable *cis*-Pt results have been reported in *FANCA*-mutant sporadic HNSCC cell lines SCC-CAL-27 and SCC-CAL-33 by Errazquin and colleagues (21). The dose-dependent sensitizations were most reliably suppressed by *FANC* transgenes in sporadic *FANCA*-mutant HNSCC cell lines, and in the FA patient-derived cell line VU-SCC-1365 (Figure S5A,B).

The kinase inhibitors gefitinib and afatinib, reported to modify the growth of FA-deficient HNSCC cell lines in culture and in xenograft experiments (40), were growth suppressive in two FA patient-derived and one sporadic *FANCA*-deficient HNSCC cell line. However, we saw little change in dose-dependent growth inhibition following *FANC* transgene complementation (Figure S5C,D). Similar results were observed for the WEE-1 kinase inhibitor AZD-1775 (Figure S5E). We saw comparable results when treating with the ATR inhibitor VE-822, and with metformin and rapamycin, two small molecules being explored to prevent, delay or treat bone marrow failure or head and neck cancer in FA (additional results not shown).

### Generation of cell line-derived 3D ‘tumoroids***’***

Many HNSCC cell lines are able to form mouse xenografts, and tumor tissue as well as some HNSCC cell lines have been used to generate HNSCC-derived organoid cultures (40-42). We used two isogenic HNSCC cell line pairs, from FA patient-derived VU-SCC-1131 and sporadic JHU-SCC-FaDu and modification of a previously reported protocol (30) to generate three-dimensional (3D) FA-proficient and -deficient *in vitro* ‘tumoroids’. Both isogenic cell line pairs readily formed tumoroids after single cell seeding in Matrigel, followed by gentle mechanical dissociation to allow subsequent growth in suspension. Tumoroid cell numbers and sizes were loosely correlated with cell line proliferation rates, time in culture and the extent of central necrosis (Figure 6).

**Figure 6.**
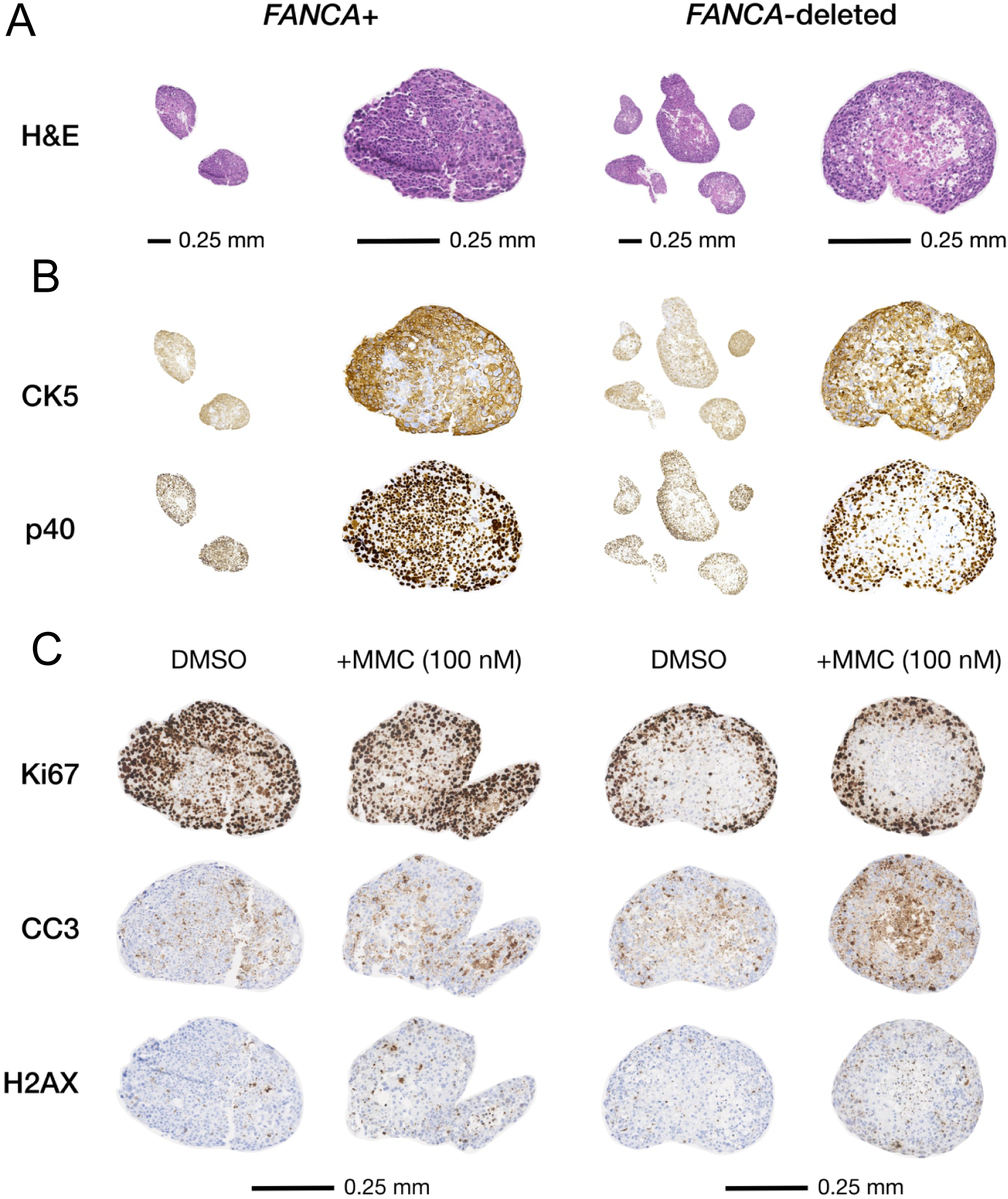
*FANCA+/FANCA*-mutant HNSCC ‘tumoroids’ generated from isogenic JHU-SCC-FaDu cells. **A**, hematoxylin and eosin (H&E) staining reveals viable squamoid cells with pleomorphic nuclei, plentiful cytoplasm, and conspicuous mitotic figures surrounding areas of variable central necrosis (4X original magnification left, 16X original magnification right). **B,** *FANCA*+ and *FANCA*-mutant tumoroids display squamous/epidermoid differentiation with diffuse nuclear p40 and cytoplasmic cytokeratin 5 (CK5) staining (original magnification 16X). **C,** Cell proliferation revealed by Ki67 immunostaining, cell death by cleaved caspase 3 (CC3) staining, and DNA damage response induction by staining for γ-H2AX (H2AX) induction, all after 3 days of continuous MMC treatment of 40 day tumoroids in suspension (original magnification 16X). All scale bars are 0.25 mm.

JHU-SCC-FaDu-derived tumoroids displayed diffuse squamous differentiation irrespective of *FANC* genotype, with strong cytokeratin 5 cytoplasmic (CK5) and p40 antigen nuclear staining (Figure 6B). Continuous cell proliferation and death were observed up to 6 weeks in culture, as evidenced by robust Ki67 and cleaved caspase 3 (CC3) immunostaining (Figure 6C). Both parental and *FANCA*-deleted JHU-SCC-FaDu Day 40 tumoroids were MMC-responsive, as indicated by CC3 and γH2AX immunostaining after 3 days of continuous MMC treatment (Figure 6C). These experiments establish the feasibility of using *in vitro* tumoroids to further explore growth, gene expression and therapeutic response profiling as a function of *FANC* genotype.

## Discussion

Fanconi anemia confers a remarkably high risk of developing carcinomas in the squamous mucosal epithelium lining the upper aero-digestive tract (oropharynx and proximal esophagus), vulva and anus (3,43,44). Skin also appears to be at an elevated, though lesser, risk of developing squamous carcinomas (4). A better understanding of cancer risk in FA - and better ways to investigate the origins, treatment, and prevention of these cancers -has been hampered by the lack of tractable, well-characterized disease models. The Fanconi Anemia Cancer Cell Line Resource (FA-CCLR) was developed to address this need by generating and distributing well-characterized *FANC*-isogenic cell line pairs of FA patient-derived or sporadic HNSCC cell lines (Figure 1).

Isogenic FA patient-derived cell line pairs were developed for three FA complementation groups that encompass ∼75% of all clinically-ascertained FA patients (2). We also generated new *FANCA*-mutant sporadic HNSCC cell line pairs for comparison with *FANC*-mutant, FA patient-derived cell lines by Cas9-targeted inactivation followed by *FANCA* transgene complementation. *FANCA* was chosen to inactivate as it is mutated in approximately two thirds of FA patients; it encodes a central protein in the heteromeric FA ‘core complex’ that orchestrates FA-dependent DNA damage responses including FANCD2 monoubiquitylation (36); and it was altered in three of our five FA patient-derived HNSCC cell lines (Table 1).

Isogenic derivatives of patient-derived or newly generated *FANCA*-mutant sporadic HNSCC cell lines were generated by *FANC* transgene transduction and/or chromosome 4 safe harbor site insertion. Both approaches restored FANC protein expression, though only viral transgene expression with higher levels of transgene protein expression fully restored MMC resistance and other FA-associated cellular phenotypes in FA patient-derived HNSCC cell lines. The requirement for higher levels of transgene protein expression to suppress MMC sensitivity in FA patient-derived cell lines may reflect the many additional genomic and other differences between FA-patient-derived and newly generated, *FANCA*-mutant sporadic HNSCC cell lines (9,12). The decision to distribute complemented pools of cells was informed by early experiments that revealed little evidence for cell line clonal heterogeneity in either FA patient-derived or sporadic HNSCC cell lines.

We observed considerable intrinsic heterogeneity across cell lines as a function of origin and genotype. However, apart from MMC and *cis*-platinum, we did not see substantial *FANC* genotype- and dose-specific differences in drug or small molecule-mediated dose-dependent survivals among *FANC*-isogenic cell line pairs. This heterogeneity was anticipated in light of the highly variable drug response profiling data for HNSCC and other cancer types made available as part of the CCLE (10,45); GDSC (46–48); and DepMap (depmap.org/portal/)) initiatives. Oxaliplatin may be the most interesting of the additional drugs tested. Despite displaying little *FANC* genotype- and dose-dependent cell killing, oxaliplatin has shown little cross-resistance with *cis*-Pt in HNSCC cell lines (49) and displays the ability to potentiate anti-PD-1 immunotherapy in preclinical models (50). These differences may reflect a different mechanism of action for oxaliplatin versus other platinum-based therapeutics such as *cis-* and carbo-Pt and potential utility as a HNSCC therapeutic agent (51,52).

This panel of isogenic FA patient-derived and newly *FANCA*-deficient HNSCC cell line pairs should provide a well-defined resource for identifying and exploring FA-specific, versus more general, aspects of HNSCC genomic instability, cellular phenotype, and therapeutic response. All of the isogenic cell line pairs completed for distribution were confirmed to display *FANC* genotype-dependent FA cellular phenotypes, and were re-authenticated and verified to be *Mycoplasma*-free prior to distribution.

Cell lines continue to play a central role in cancer research (53–55). They are experimentally tractable disease models that can capture and propagate many cell-autonomous features of neoplasia, and by virtue of simplicity and low cost have enabled many high-throughput drug, small molecule and genomic screening protocols among other experimental analyses. The resulting publicly available data have, in turn, become a widely used resource and cancer discovery tool (see, e.g., CCLE (10,45); GDSC (46–48); DepMap resources (depmap.org/portal/)).

The limitations of cancer cell line models are also well-recognized, most notably the lack of immune/inflammatory cells and a vascularized tumor microenvironment. These limitations are now being effectively addressed to pursue specific questions of tumor biology or therapeutic response by combining xenograft, tissue engineering, and organoid protocols. The *FANC*-isogenic cell line pairs generated as part of the FA-CCLE can be used in many of these approaches, and can provide highly reproducible results once cell line-intrinsic differences are taken into account and care is taken to minimize passage-associated phenotypic drift or genomic evolution (53,55,56). Our hope is that the FA-CCLR will support many promising new research directions to benefit individuals with FA and their families as well as individuals with sporadic HNSCC.

## Acknowledgements

We thank members of the FA research community for locating and providing FA cancer cell lines, and resolving questions of cell line origins or identity. Hui Li assembled our initial *FANC* gene open reading frame vectors, Helmut Hanenberg provided detailed information on retroviral complementation vector structure and Bryan Johnson of the UW-ISCRM Histology and Imaging Core provided expert guidance for tumoroid histologic analyses. We dedicate this manuscript to the FA community of patients, families, and scientists, and to our late colleague Eddie Mendez (Fred Hutchinson Cancer Research Center, Seattle, WA USA). Eddie was an inspiring colleague and physician-scientist dedicated to helping patients and families facing HNSCC and FA. Many of the ideas for a FA-CCLR were first discussed with Eddie.

## Supplemental Data

### 1. Extended Methods

#### Cell line sources

Twelve FA patient-derived or sporadic HNSCC cell lines, identified from reports or via personal communication, were acquired and given systematic names in accordance with ICLAC (International Cell Line Authentication Committee) guidelines (1,2) to avoid ambiguity or confusion (Table 1). Cell line identities were established by short tandem repeat (STR) DNA marker profiling using the CellCheck9 Plus panel of 9 human species-specific STR markers (IDEXX BioAnalytics, Westbrook, ME USA). Profiles were compared with previously published STR data where they existed, or reported here to provide cell line-specific identification and authentication signatures (Table S1). The same assay panel included markers to identify interspecies contamination by mouse, rat, African green monkey or Chinese hamster cells, and to detect the presence of *Mycoplasma* ssp. infection.

FA-patient-derived cells CCH-SCC-FA1 and CCH-SCC-FA2 were established from de-identified tumor samples from male (FA1) and female (FA2) FA-patients, obtained through the National Disease Research Interchange (NDRI) by Susanne Wells (Cincinnati Children’s Hospital, Cincinnati OH). CCH-SCC-FA1 was initiated from a tongue lesion, and CCH-SCC-FA2 from an oral cavity lesion (3,4). Both were grown initially on irradiated NIH-3T3 mouse feeder cells, from which feeder-independent sublines were developed by Susanne Wells and provided to Josephine Dorsman (Amsterdam University Medical Center, Amsterdam, the Netherlands) who contributed the cultures used to develop FA-CCLR isogenic pairs (5).

The FA patient-derived cell line OHSU-SCC-974 was isolated from a biopsy of an oropharyngeal lesion in a 29 y.o. male FA patient of the *FANCA* complementation group (6). The initial cell line was provided by Grover Bagby and Laura Hays (OHSU, Portland, OR) to Susanne Wells (Cincinnati Children’s Hospital Medical Center, Cincinnati, OH). A *FANCA*-complemented isogenic cell line pair was developed from a genome-sequenced culture provided by Agata Smogorzewska (Rockefeller University, New York, NY).

FA patient-derived HNSCC cell lines VU-SCC-1131 and VU-SCC-1365 were established from biopsies of oral mucosal lesions in a 34 year old *FANCC*-mutant female (VU-SCC-1131) and a 25 year old *FANCA*-mutant male (VU-SCC-1365)(6). The FA patient-derived HNSCC cell line VU-SCC-1604 was established from a biopsy of a tongue lesion in a *FANCL-*mutant female FA patient (3). All three VU cell lines were provided by Josephine Dorsman and Ruud Brakenhoff of the Amsterdam University Medical Center, Amsterdam, the Netherlands.

Sporadic HNSCC cell line JHU-SCC-FaDu (ATCC HTB-43) was established from a punch biopsy of a hypopharyngeal tumor (7), and SFCI-SCC-9 (ATCC CRL-1629) from a biopsy of a tongue SCC in a 25 year old male (8). Both were purchased from the American Type Culture Collection (Manassas, VA, USA), with an agreement from ATCC dated 4 January 2022 to allow derivatives of these cell lines to be distributed without additional cost upon approval of the transfer from FA-CCLR to ATCC-approved end users. See the ATCC Transfer Request website for additional information on transfers: https://transferrequest.atcc.org/trifform.aspx). ATCC requested that parental JHU-SCC-FaDu (ATCC HTB-43) and SFCI-SCC-9 (ATCC CRL-1629) cell lines be purchased directly from ATCC.

Sporadic HNSCC cell lines UM-SCC-1 and UM-SCC-47 were originally established by Thomas Carey and colleagues (9,10) and Randall J. Kimple (11). A culture of UM-SCC1 was provided by Susanne Wells (Cincinnati Children’s Hospital Medical Center, Cincinnati, OH), and of UM-SCC-47 by the late Eddie Mendez (Fred Hutchinson Cancer Research Center, Seattle, WA USA). The reporting of data generated using UM-SCC-47 is reported here with permission of Thomas Carey, University of Michigan, Ann Arbor MI), and granted on 8 December 2021.

Sporadic HNSCC cell lines CAL-SCC-27 and CAL-SCC-33 were derived from tongue lesions in a 56 year old male (CAL-SCC-27) and a 69 year old male (CAL-SCC-33)(12–14). *FANCA-* mutant and *FANCA* retroviral transgene-complemented derivatives for each cell line were developed and provided by Ricardo Errazquin and Ramon Garcia-Escudero (CIEMAT, Madrid Spain) as part of assembling the FA-CCLR Resource (15).

The human osteosarcoma cell line cU-2 OS (ATCC HTB-96), established from a moderately differentiated sarcoma in the tibia of a 15 year-old girl (16), was purchased from the American Type Culture Collection (Manassas, VA, USA) and used for reagent testing and protocol development.

#### Cell culture and cell line authentication

FA-patient derived OHSU-SCC-974 and sporadic JHU-SCC-FaDu HNSCC cell lines were grown in Eagle’s Minimum Essential Media (MEM) supplemented with 10% fetal bovine serum (FBS, Hyclone Laboratories, South Logan Utah) and 1% penicillin-streptomycin (Gibco #10378016). Cell lines UM-SCC-1, UM-SCC-47, CCH-SCC-FA1 and FA2, VU-SCC-1131, VU-SCC-1365, VU-SCC-1604, and U-2 OS were grown in Dulbecco-Modified MEM (D-MEM) supplemented with 10% FBS, 1% non-essential amino acids (Gibco #11140050) and 1% penicillin-streptomycin. Sporadic HNSCC cell lines SCC-CAL-27 and SCC-CAL-33 do not require 1% non-essential amino acids for growth (15). SFCI-SCC-9 cells were grown in a 1:1 mixture of D-MEM/Nutrient Mixture F12, supplemented with 10% FBS, 0.5 mM sodium pyruvate, 400 ng/mL hydrocortisone and 1% penicillin-streptomycin. All cell lines were grown at 37°C in a 5% CO_2_/ambient (20%) oxygen, water vapor-saturated incubator.

#### Cell line genomic characterization

Prior reports and more recent, comprehensive genomic profiling data were combined and used to identify cell line-associated single nucleotide variants (SNV), copy number variants (CNV) and cell line SNV/indel variant (17–21). Somatic SNV/indel and copy-number data for FA patient-derived cell lines OHSU-SCC-974, CCH-SCC-FA1 and CCH-SCC-FA2 were extracted from Webster et al. (20). In brief, somatic SNV/indel calls were made using CaVEMAN (22) and Pindel (23), using tumor cell line and patient matched control whole-genome sequencing (WGS) data. Copy-number amplifications and deletions in OHSU-SCC-974, CCH-SCC-FA1 and CCH-SCC-FA2 were called using CNVkit (24) to compare tumor cell line and patient-matched WGS data. SNV/indels and copy-number amplifications and deletions were called for FA patient-derived cell lines VU-SCC-1604, VU-SCC-1365 and VU-SCC-1131 using, respectively, Mutect2 (25) and CNVkit to compare tumor cell line and patient-matched fibroblast whole-exome sequencing (WES) data (21). In light of restricted primary data access for these three lines, we focused on variant calling in genes previously shown to be significantly mutated in HNSCC, and considered to be potential cancer drivers when altered (26,27). None of these 35 HNSCC-associated genes was mutated in U-2 OS, which is *TP53*+/*FANC*+ (18,28).

For sporadic HNSCC cell lines JHU-SCC-FaDu, CAL-SCC-27, CAL-SCC-33, SFCI-SCC-09 and UM-SCC-01, somatic mutation data was extracted from the Broad Institute Cancer Cell Line Encyclopedia (CCLE)(18), employing Mutect2 and Absolute (29) for SNV/indel and copy-number analyses. For all cell lines, the reporting threshold for copy-number amplifications and deletions was set at log2(CN)>0.5 or log2(CN)<-0.5 respectively.

#### Generation of *FANCA-*mutant sublines of sporadic HNSCC cells

The systematic workflow used to generate *FANC*-isogenic cell line pairs from sporadic HNSCC is shown in Figure 1. Biallelic inactivation of the endogenous *FANCA* genes in sporadic HNSCC cell lines and in control U-2 OS cells was generated by dual guide RNA-Cas9 mediated deletion of a 5’ portion of the FANCA gene to disrupt the intron 1 splice donor site, exon 2, and a portion of exon 3 (Figure 3A). The expression vector pCas9-GFP-FA1&4 was constructed by replacing the dual gRNAs in plasmid pUS2-SH231 (Addgene, plasmid #115150) with pairs of gRNAs in the 5’ end of the *FANCA* gene that were identified using CHOPCHOPv3 (30). Different *FANCA* gene-targeting gRNA pairs were tested empirically to identify the best-performing guides as assessed by deletion efficiency.

In brief, cell lines were transfected with pCas9-GFP-FA1&4 plasmid vectors encoding pairs of gRNAs using either polyethylenimine (PEI) or electroporation. Sterile flow sorting was used to enrich transfected, GFP+ cells 3 days after transfection for seeding at low density (500 cells/10-cm dish) to generate single cell-derived colonies after 10-14 days additional growth with medium changes every 3-4 days. Colonies were transferred by sterile filter paper disc transfer into 48-well plate wells for expansion, the generation of replicate cultures and screening by PCR to identify colonies that had targeted deletions in one or both *FANCA* alleles. The best-performing guide RNA pair identified by this approach were FA4, targeting *FANCA* intron 1 sequence 5’ - GAACCGACTTCTCTCCGTAG - 3’ and FA1, targeting *FANCA* exon 3 sequence 5’ - GATTGACTGTGACAGTTCTG - 3’. Oligonucleotides encoding both guides were cloned into a dual expression vector to generate pCas9-GFP-FA1&4 (Figure 3A). We did not find evidence of stable integration with expression of Cas9 or CpfI after transient transfections, and observed no stable GFP+ cells. This suggests that knockout/complemented cells express little or no residual Cas9 or Cpf1 protein.

Colony PCR FANCA deletion screens were performed using amplification primers flanking the target positions of *FANCA* gRNAs in intron 1 and intron 3: the forward primer used, FA_FWD (5’ - AATTGTTCTCCCGTCTGCTCTC - 3’) was located in intron 1, and the reverse primer FA_REV (5’ - GGGCCGTCTCCGTTAGTTTC - 3’) in Intron 3 (Figure 3A).

Single cell-derived colonies were subcloned and grown to ≥ 50% confluency in 24 well plate wells, then detached using trypsin/EDTA, pelleted by centrifugation at 1000 rpm for 5 min in 1.7 ml microfuge tubes, then resuspended and lysed by the addition of 100 µl of DirectPCR Lysis Reagent containing 200 µg/ml proteinase K (Viagen, Los Angeles, CA, USA). PCR reactions were performed in 20µl reaction volumes containing 1 μl of DNA lysate (equivalent of 1000-2000 cells), 8.2 μl deionized H_2_O, 0.4 μl each of 15 μM FA_FWD and FA_REV primers, and 10μl Hot-Start PCR Ready Mix (KAPA biosystems, Catalog #KK5609). The thermocycling amplification program was: denaturation at 94°C for 6 min; 35 cycles of amplification consisting of 94°C for 20s, 56°C for 20s, and 72°C for 2 min; then final extension at 72°C for 5 min. PCR reaction products were analyzed by electrophoresis through a 1% (w/v) 6 cm agarose gel run at 100V in Tris acetate EDTA buffer containing GelGreen™nucleic acid gel stain (Genecopoeia, Catalog #N101) until the predicted DNA fragment size range could be clearly resolved versus a size standard. Gels were imaged on a FluorChem FC2 imager system (ProteinSimple - Alpha Innotech, San Jose, CA).

Errazquin *et al.* used a parallel strategy to generate *FANCA*-mutant derivatives of the sporadic HNSCC cell lines CAL-SCC-27 and CAL-SCC-33 (15). In brief, *FANCA* exon 4-targeting gRNA-Cas9 ribonucleoprotein particles were transfected into cells followed 3 days later by Surveyor™ Mutation Detection (IDT), then FACS sorting of Surveyor-positive cultures to isolate and expand single cell-derived colonies in conditioned medium. PCR amplification and Sanger sequencing were used to confirm the presence and nature of biallelic *FANCA* mutations.

#### Generation of *FANC transgene*-complemented cell line clones and pools

FA-patient derived *FANC* mutant and newly generated *FANCA-*deletion sporadic cell lines were *FANC* transgene-complemented using both chromosomal safe harbor site targeted insertion and viral transduction (Figure 1). Safe harbor site complementation used Cas9 or CpfI and site-specific gRNAs to target cognate *FANC* transgenes at the chromosome 4q Safe Harbor Site 231 (SHS231)(Figure 3C)(31). Safe harbor site targeting aimed to provide complemented cells with more uniform transgene copy numbers at a defined genomic location to use to generate *FANC*-complemented cell clones and pools. Lentiviral or retroviral complementation was used as an alternative to safe harbor site insertion to achieve higher *FANC* transgene expression levels. This was required in patient-derived FA-deficient cell lines to more fully complement MMC sensitivity and/or other FA-deficient phenotypes (see manuscript Results and Discussion for additional detail).

SHS231-specific transgene vectors were constructed for complementation groups *FANCA, FANCC* and *FANCL,* in which HA epitope-tagged *FANC* transgene linked via a T2A peptide to either a puromycin or hygromycin selection cassette was expressed under the control of an EF1α promoter (Figures 3C and S3). SHS231 safe harbor site complementation transgenes were electroporated into *FANC*-deficient cells with expression plasmids encoding Cas9 or Cpf1 and SHS231 safe harbor site-targeting gRNAs. Transfected cells were incubated for 72 hs in culture medium prior to adding hygromycin B (400μg/mL; Millipore Sigma, Catalog #400050) or puromycin (1µg/ml; Invivogen) to promote the selective outgrowth of transgene-positive colonies that were subcloned by sterile filter paper disc transfer into 48-well dishes and expanded under selection for an additional 10-14 days prior to PCR confirmation of transgene presence and orientation. PCR primer pairs were anchored in genomic DNA adjacent to SHS231 on Chr 4, and in transgene sequence adjacent to the predicted SHS231-transgene junction sequence T231_F8: 5’ - AGAACATGCAATGGCTAGC - 3’, and transgene vector reverse primer attP_R1: 5’ - GCGGTGGTTGACCAGACAAA - 3’. This primer pair yields a PCR product of 634 bp when transgenes were inserted at SHS231 in the correct orientation (Figure 3C).

Lentiviral complementation was performed using 3rd generation HIV-1-derived vectors where *FANC* transgene expression was under EF1α promoter control and linked to puromycin or hygromycin selection cassette expression. *FANC* transgenes were assembled from cDNA clones (*FANCA*) or synthesized from reference nucleotide sequences NG_007425.1 (*FANCF* NCBI Reference Sequence) and NG_007418.1 (*FANCL* NCBI Reference Sequence) by IDT (Coralville, Iowa USA) as oligonucleotides. The resulting transgene cassettes were then assembled by Gibson cloning and sequenced to confirm their structure and sequence integrity. Large stocks of completed vectors were grown in *E.coli* and purified by CsCl gradient centrifugation for packaging (see: https://www.addgene.org/protocols/plko/#E for the detailed protocol). Lentiviral complementations were performed at low multiplicities of infection (MOI) as assessed by cell fraction surviving after selection, to ensure low mean viral copy numbers. The retroviral vector used to complement *FANCA*-mutant sublines of CAL-SCC-27 and CAL-SCC-33 was S11FAIN *FANCA,* in which transgene expression was coupled via an IRES to a neomycin/G418 resistance cassette. This vector has been described in (15), with additional detail provided by Helmut Hanenberg, University Hospital Essen Germany (personal communication October 2021)(Figure S3). S11FAIN *FANCA* was propagated, packaged and used in transductions as previously described (32).

#### Cell proliferation assays

Proliferation rate analyses were used to identify potential metabolic alterations or constraints as a function of cell line genotype and growth medium. Cells were seeded (5,000/well) in 48- well plates in D-MEM, MEM, or RPMI 1640 base medium supplemented with 10% FBS and penicillin-streptomycin (Day 0), then manually counted 24 hr later (Day 1) and again after 3 additional days (on Day 4) of undisturbed growth at 37°C in water vapor-saturated incubators in a 5% CO_2_, 20% oxygen atmosphere. Doubling day proliferation rates were calculated as previously described to reflect the 4 day growth interval (33):

> Proliferation rate = log_2_[final cell count (Day 4)/initial cell count (Day 1)]/3.

#### Cell cycle analyses

Cell cycle phase distribution profiles were determined for isogenic cell line pairs using exponentially growing control and mitomycin-C-treated cultures. In brief, 2.0 × 10^5^ exponentially growing cells were plated on 60 mm dishes in medium without or containing 10 nM MMC for 72 h, prior to centrifugation and resuspension in 1x DAPI solution (146 mM NaCl, 10 mM Tris pH 7.4, 2 mM CaCl_2_, 22 mM MgCl_2_, 0.1 mg/mL BSA, 0.1% IGEPAL, 10 μg/mL DAPI, 10% DMSO) at 2.0 × 10^6^ cells/ml, followed by trituration through a 25 gauge needle to release intact nuclei. Stained nuclei were analyzed immediately, or stored at -20 °C until flow cytometric profiling for DNA content on a BD Biosciences LSR II Flow Cytometer versus a stained chicken erythrocyte DNA standard as previously described (34). Flow output analysis was performed using FlowJo flow cytometry data analysis software (BD Biosciences).

#### Drug and small molecule response profiling

The ability of drugs or small molecules to suppress cell proliferation as a function of dose was determined for MMC and for nine other drugs or small molecules by continuous growth for 4 days without refeeding, followed by WST-1 (Takara Bio, Catalog #MK400) or alamarBlue (ThermoFisher, Catalog #DAL1025) staining to compare cell numbers in control and treated cultures. In addition to MMC, we tested *cis*-Pt, carboplatin and oxaliplatin; formaldehyde; the kinase inhibitors gefitinib and afatinib; the WEE-1 checkpoint kinase inhibitor AZD-1775; M6620 (VX-970), an ATR inhibitor now entering clinical trials; and the metabolic inhibitors metformin and rapamycin. In brief, 2,500 cells/well were plated in triplicate in 96-well plate format in 200 μl of complete growth medium followed by the addition of a two-fold serial dilution of mitomycin-C, other drug or small molecule or control medium (Millipore Sigma Catalog #475820). Data were normalized versus control medium wells lacking drug (Figures 4, S4 and S5, Table S2).

#### Protein detection and characterization by Western blot analyses

Western blot analyses were performed using total cellular protein extracts prepared from ∼ 2.5×10^6^ cells in RIPA Lysis Buffer (Boster Biological Technology, Pleasanton CA, Catalog # AR0105) as previously described (35). Protein concentrations were determined using the Pierce BCA Protein Assay Kit (Thermo Fisher, Catalog #23227), and 40 μg of protein lysate/lane were loaded and size-fractionated by electrophoresis through precast 4-12% Bis-Tris protein gels (Thermo Fisher Catalog #NP0336BOX). Proteins were transferred onto nitrocellulose membrane (Thermo Fisher Catalog #88025), then blocked in Tris-buffered saline solution containing 5% non-fat dry milk (w/v) for 1 hr at room temperature or overnight at 4°C prior to incubation with primary antibodies to detect endogenous or transgene-encoded proteins. In order to detect FANCA, membranes were incubated with a rabbit FANCA-primary antibody (Bethyl A301-980A, Bethyl Laboratories TX, USA) followed by detection with an HRP-conjugated anti-rabbit IgG goat polyclonal antibody (Cell Signaling, cat #:7074) at 1:1500 dilution. FANCC, FANCF and FANCL proteins were detected using the same general protocol and an anti-FANCC antibody at 1:1000 dilution (MA5-16138, Pierce Biotechnology, MA, USA); an anti-FANCF antibody at 1:500 dilution (sc-271952, Santa Cruz Biotechnology, TX, USA); or an anti-FANCL antibody at 1:500 dilution (sc-137067, Santa Cruz Biotechnology, TX, USA). All *FANC* transgene-encoded except FANCA could also be detected by immunoblotting for an N-terminal HA epitope tag using the anti-HA-tag-directed mouse monoclonal antibody 901501 (Biolegend, CA, USA) at 1:1000 dilution.

#### Protein detection and characterization by targeted mass spectrometry analyses

Peptide immunoenrichment coupled with targeted multiple reaction monitoring mass spectrometry (immuno-MRM) was used as a complementary approach to detect and quantify the expression and post-translational modification status of FANCD2 and additional DNA damage response and repair (DDR) proteins prior to and after MMC treatment. The two immuno-MRM panels used for these analyses target 126 peptides and 62 post-translational modification sites in 66 DDR proteins including 10 FANC proteins (36–38).

In brief, immuno-MRM assays were performed using isogenic pairs of *FANCA*-deleted or control JHU-SCC-FaDu HNSCC or U-2 OS osteosarcoma cells seeded at 5 × 10^6^ (FaDu pair) or 2 × 10^6^ (U-2 OS pair) cells/10-cm dish in triplicate, then grown for 24 hr in the presence or absence of 0.2µM mitomycin C. Cells were then harvested to prepare whole cell lysates as previously described (36,37). Lysates were reduced, alkylated with iodoacetamide, and digested with Lys-C (Wako, #129-02541) at a 1:50 enzyme:protein mass ratio, followed 2 hr later by the addition of trypsin (#V5111, Promega, Madison, WI) at a 1:100 enzyme:protein ratio followed by overnight incubation at 37°C with shaking. Stable isotope-labeled peptide standards (New England Peptide, Gardner, MA) were then added at 150 fmol/mg prior to desalting. Seven peptide standards were added at four-fold high concentration (to 600 fmol/mg each) to compensate for lower relative MS signal levels on the mass spectrometer: CHEK1:YSSSQPEPR; CHEK1:YSSpSQPEPR: MDC1: AQPFGFIDpSDTDAEEER; CLU: ASSIIDELFQDR; PCNA: AEDNADTDLALVFEAPNQEK: RAD50: LFDVCGSQDFESDLDR and RAD50: LFDVCGpSDFESDLDR. Antibodies were crosslinked on protein G beads (GE Sepharose, Cat. # 28-9513-79), and the flow-through from enrichment of the first antibody panel was used as input for the second antibody panel purification. All eluates were frozen and analyzed independently by mass spectrometry as previously described (36–38), using independent runs for the enriched peptides from the two panels.

LC-MRM-MS was performed on an Eksigent Ultra nanoLC system with a nano autosampler and chipFLEX system (Eksigent Technologies, Dublin, CA), coupled to a 5500 QTRAP mass spectrometer (SCIEX, Foster City, CA) operated in positive ion multiple reaction monitoring (MRM) mode. Peptides were loaded onto a trap column (Reprosil C18, 5 mm × 200 m) at 5 L/min for 3 min using mobile phase A (0.1% formic acid in water). The LC gradient was delivered at 300 nL/min, and consisted of a linear gradient of mobile phase B (90% acetonitrile and 0.1% formic acid in water) from 3-14% B over 1 min, 14-34% B over 20 min and 34-90% B over 2 min followed by re-equilibration at 3% B on a 15 cm × 75 m chip column (Reprosil AQ C18 particles, 3 m; Dr. Maisch, Germany). Parameters for collision energy (CE) were taken from prior optimization of synthetic peptides (36,38). Scheduled MRM transitions used a retention time window of 150 sec and a cycle time of 1.0 sec to enable sufficient point sampling across peaks to allow MRM data to be quantified using Skyline (39)(see: https://skyline.ms/project/home/software/skyline/begin.view).

Peak integrations were reviewed manually, and transitions from analyte peptides were confirmed by alignments of the retention times of endogenous and added, heavy stable isotope-labeled reference peptides with equivalent relative recorded transition areas. Transitions with detected interferences were not used in data analyses. Integrated raw peak areas were exported from Skyline, and total intensities were calculated using Peak Area + Background. Peak areas were filtered using experimentally determined lower limits of quantification (LOQ) from response curves generated using cell lysate as the matrix. Peak area ratios were obtained by dividing light peptide peak areas by the corresponding heavy mass-labeled reference peptide. The mean and standard deviations of peak area ratios were calculated from three complete biological replicates of cell lines ± treatment, with reporting of the results from both immunopurification panels as a single list of analytes.

#### Three dimensional (3D) cell line-derived ‘tumoroids’

A previously described 3D culture protocol (40-42) was modified and used to generate 3D ‘tumoroid’ cultures from isogenic *FANC*-mutant and -complemented derivatives of sporadic HNSCC cell line JHU-SCC-FaDu, and the corresponding isogenic FA patient-derived VU-SCC-1131 cell line. In brief, exponentially growing single cells were suspended in Matrigel (Corning Life Sciences) at a ratio of 2:1 (v/v) Matrigel:cell suspension, then 40 uL aliquots containing ∼1000 cells were pipetted onto pre-warmed 24-well plates to form ‘domes’ to polymerize at room temperature for 30 min followed by curing for 1 hr at 37°C. Plates were then flooded with complete culture medium (D-MEM supplemented with 10% FBS, 0.4 μg/mL hydrocortisone (Stemcell Technologies #07925), 1% penicillin-streptomycin, 1X insulin-transferrin-selenium (Thermo Fisher #41400045), 10ng/mL human epidermal growth factor (Sigma #E9644) and 1X B27 (Gibco #17504044). After 7 d, domes were gently mechanically dissociated to free individual tumoroids for additional growth in suspension on non-adherent plates.

The mitomycin-C responsiveness of 40 d-old tumoroid cultures was assessed by growth for 3 days in suspension medium supplemented with 10, 50 or 100 nM MMC. Treated tumoroids were collected by unit gravity sedimentation, gently washed twice by resuspension in PBS followed by room temperature fixation for 24 hr in neutral-buffered formaldehyde. Fixed tumoroids were washed twice and stored at 4°C in 70% ethanol for 24 h prior to re-suspension in agarose to facilitate handling and embedding for sectioning.

Paraffin sections were stained with hematoxylin and eosin (H&E), and for differentiation/ disease-specific antigens CK5 cytokeratin 5 (CK5) and the p40 fragment of p63, as well as the functional state markers Ki67, H2AX and cleaved caspase 3 (CC3). Antibodies used for staining were: for cytokeratin 5 (CK5) Cell Marque EP1601Y antibody at a 1:50 dilution; for the p40 fragment of p63, BioCare Medical BC-28 antibody at 1:200 dilution; for the proliferation marker Ki67, Dako MIB-1 antibody at 1:100 dilution; for the DNA damage-responsive marker Ser139-H2AX phosphorylation, BD Pharmingen 560447 antibody at 1:400 dilution; and for the cell death, cleaved caspase 3 (CC3) BioCare Medical CP229B antibody at 1:250 dilution.

Antigen retrieval was attempted by heat treatment, citrate or EDTA to identify and optimize retrieval and staining for each antigen on a Leica BOND-III automated stainer (Leica Biosystems, Buffalo Grove, IL, USA).

---

**Figure S1.**
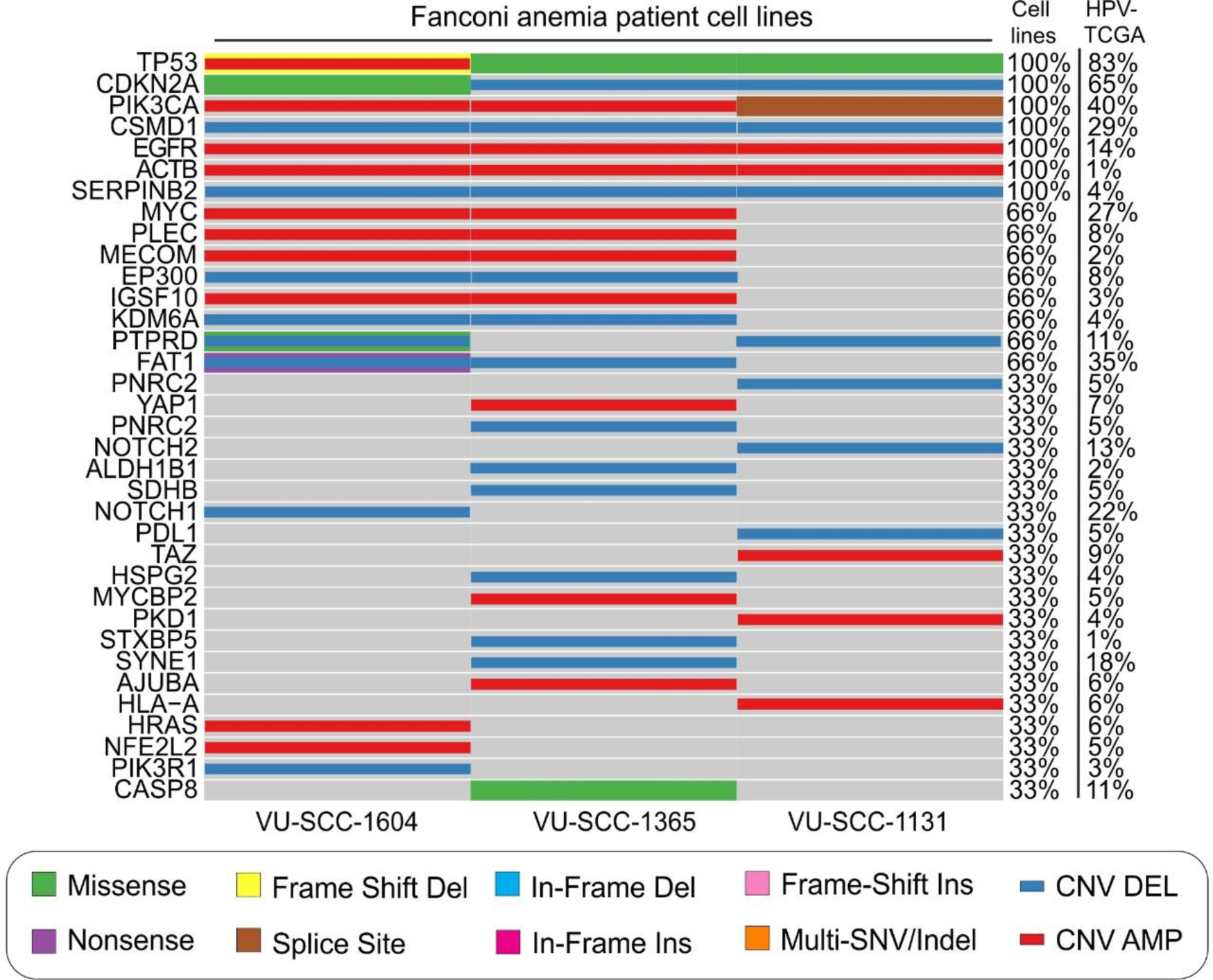
– Mutational landscape of additional Fanconi anemia patient SCC cell lines. Oncoplot of three Fanconi patient-derived HNSCC cell lines VU-SCC-1604, VUSCC-1365 and VU-SCC1131. Displayed along the left margin is a curated panel of oncogenes and tumor suppressor genes frequently mutated in SCC. The Figure key at bottom indicates the type of mutation identified, while the two columns along the right margin indicate the frequency with which each gene was altered in VU SCC cell lines versus a comparison dataset of HPV-negative HNSCC included in TCGA. The reporting thresholds for copy-number amplifications and deletions are log2(CN)>0.5 or log2(CN)<-0.5 respectively.

**Figure S2.**
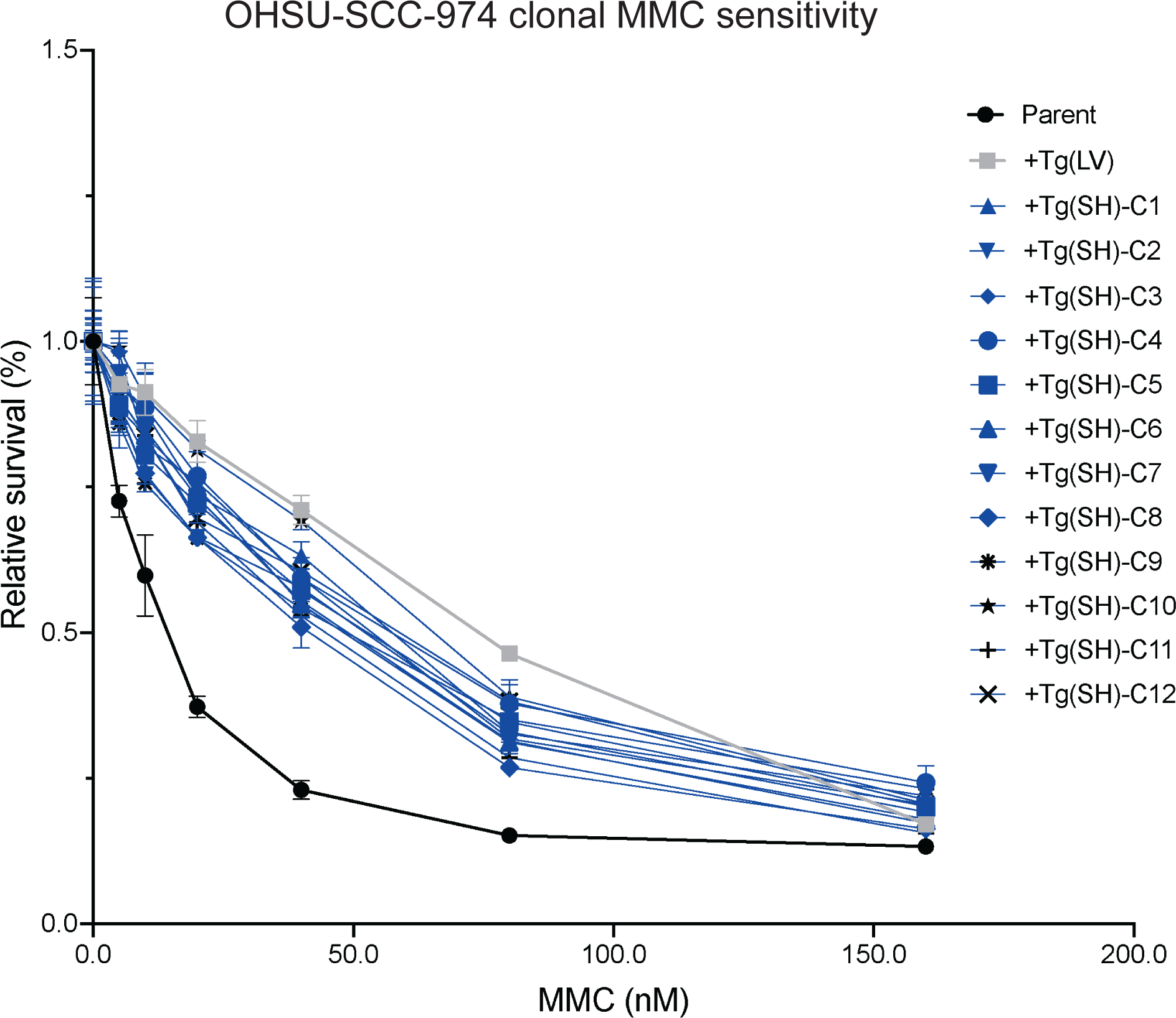
MMC dose-dependent survival following *FANCA* transgene complementation of *FANCA*-mutant, patient-derived HNSCC cell line OHSU-SCC-974. Blue lines show MMC dose-dependent survival of 12 independent, single cell-derived chromosome 4q safe harbor site *FANCA* trans-gene-complemented clones, compared with the *FANCA*-mutant parental cell line (parent, ●) and with a pool of parental cells that had been lentivirally *FANCA*-complemented (+TgLV, ▪). Data represents mean ± SEM.

**Figure S3.**
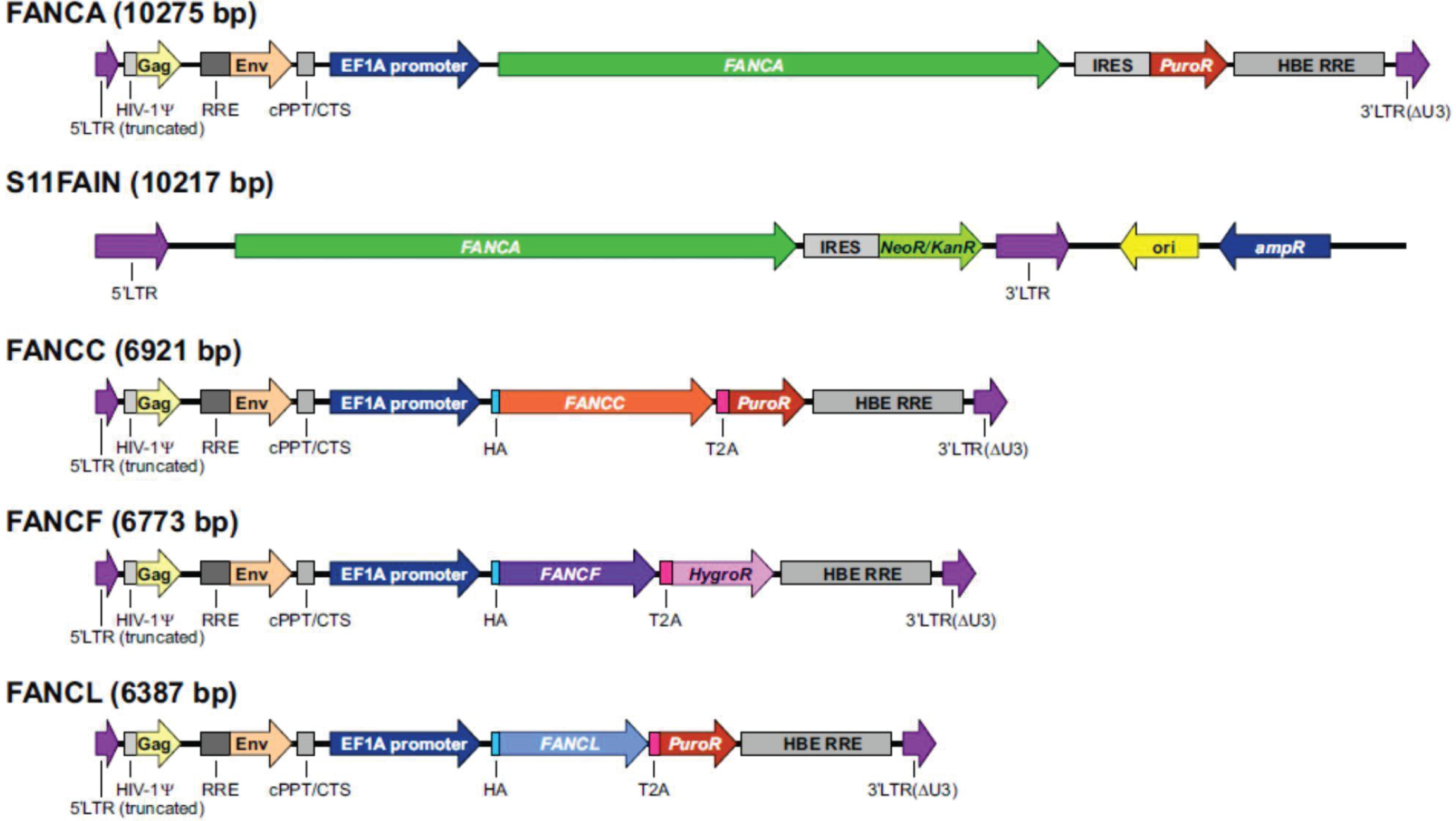
*FANC* transgene lentiviral and retroviral vectors. Structure of lentiviral vectors and the S11FAIN retroviral vector used to complement *FANC*-mutant FA patient-derived or newly generated sporadic HNSCC cell lines (see Table 1 for detail). *Genetic element abbreviations:* LTR: viral long terminal repeats; HIV-1 Ψ: packaging signal of human immunodeficiency virus type 1; RRE: Rev response element (RRE) of HIV-1 that promotes Rev-dependent RNA export from the nucleus; cPPT/CTS: central polypurine tract and central termination sequence of HIV-1; Gag, Env: HIV gag and env genes; IRES, internal ribosome entry site; HA: HA epitope tag; T2A: self-cleaving peptide spacer; NeoR/KanR, PuroR, HygroR: antibiotic selection cassettes.

**Figure S4.**
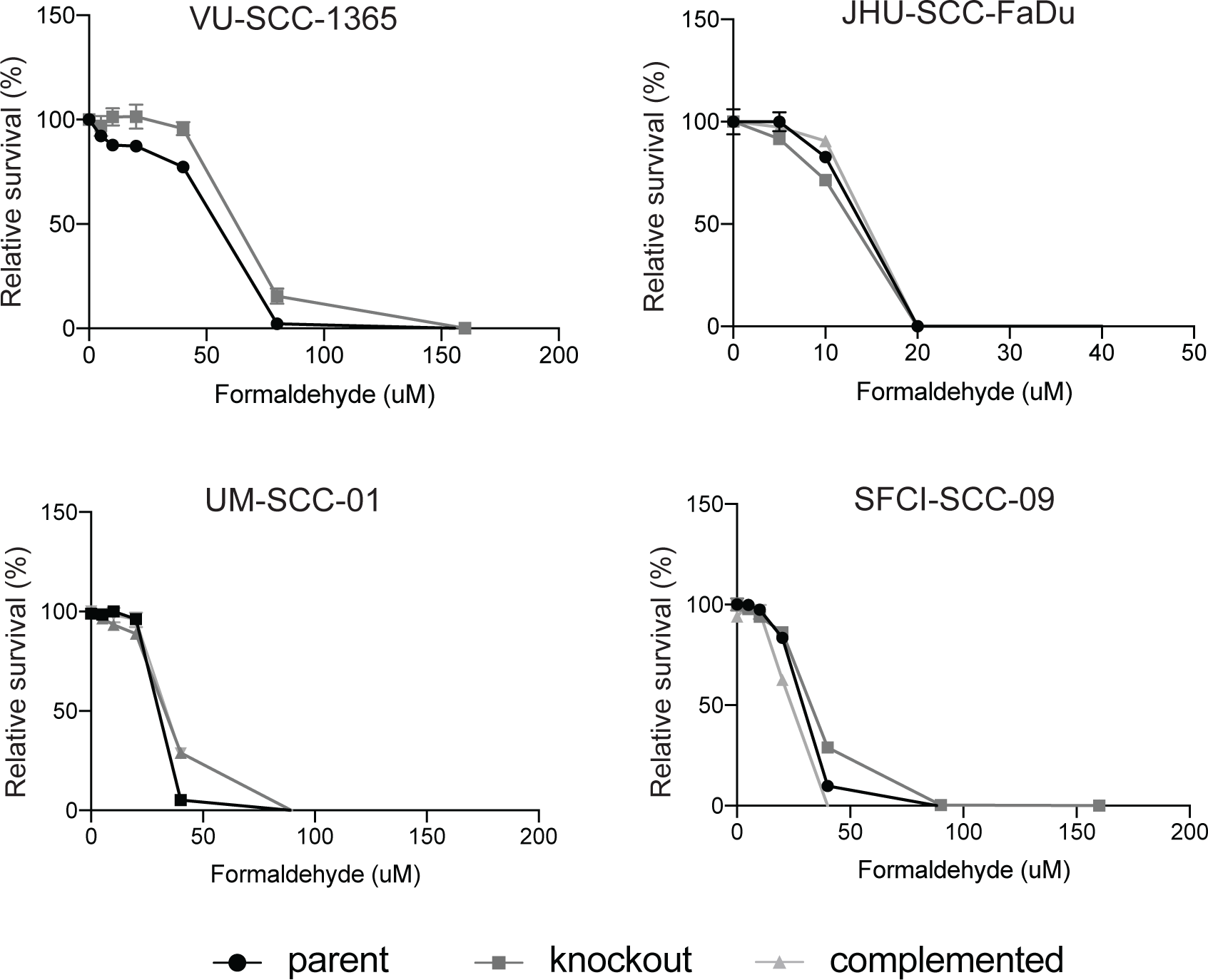
Formaldehyde sensitivity of FA patient-derived and sporadic HNSCC isogenic cell line pairs. Dose-dependent suppression of cell proliferation of *FANCA*-isogenic FA patient-derived (VU-SCC-1365) and sporadic (JHU-SCC-FaDu, UM-SCC-01, and SFCI-SCC-09) HNSCC cell line pairs to formaldehyde. Data represents mean ± SEM.

**Figure S5.**
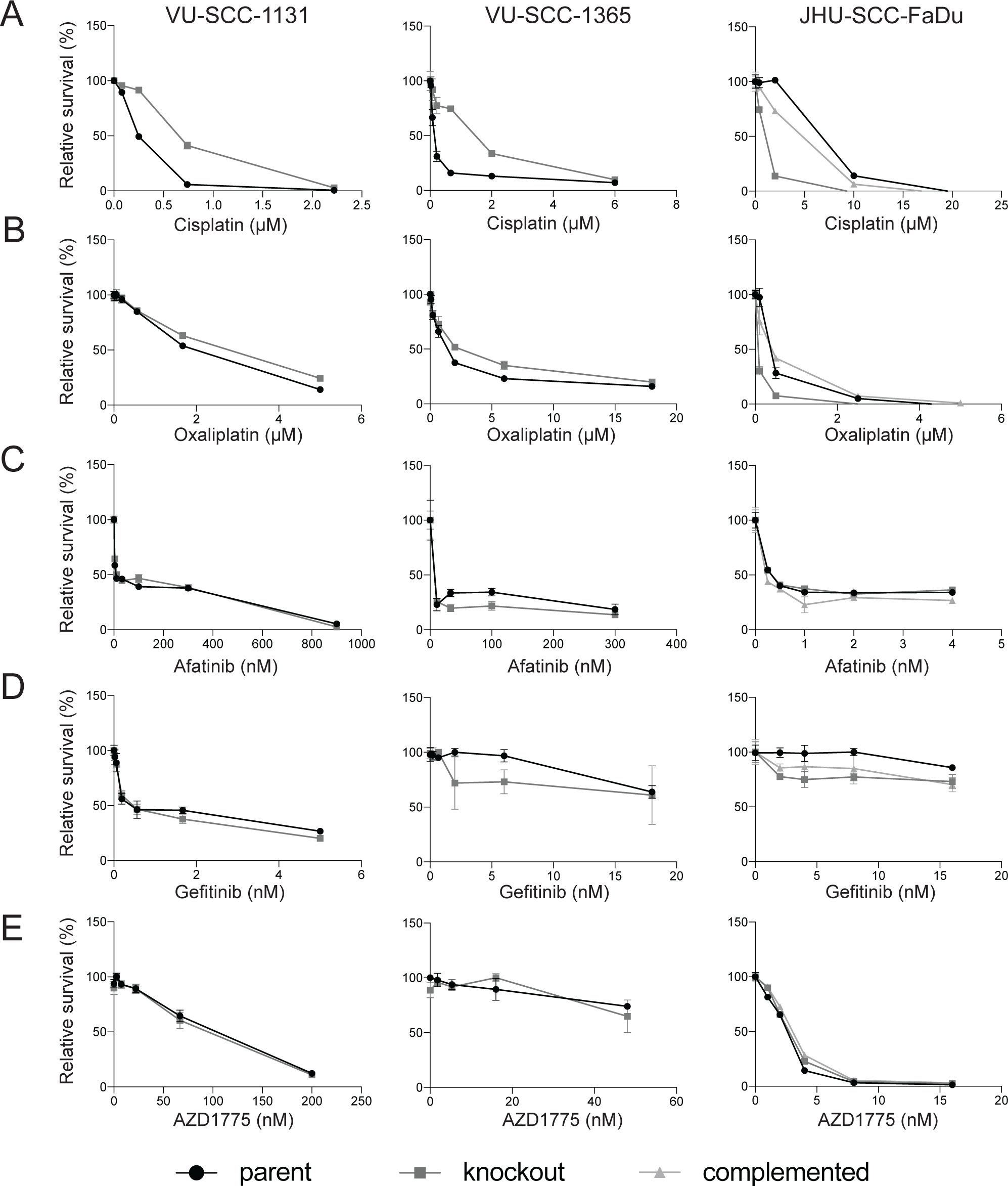
Drug and small molecule response profiling of FANC-isogenic FA patient-derived and sporadic HNSCC cell line pairs. Dose-dependent survival of *FANCC*-isogenic FA patient-derived (VU-SCC-1131), *FANCA*-isogenic FA patient-derived (VU-SCC-1365), and sporadic (JHU-SCC-FaDu) HNSCC cell line pairs to cisplatin (**A**), oxaliplatin (**B**), afatinib (**C**), gefitinib (**D**), and AZD1775 (**E**). Data represents mean ± SEM.

**Table S1.**
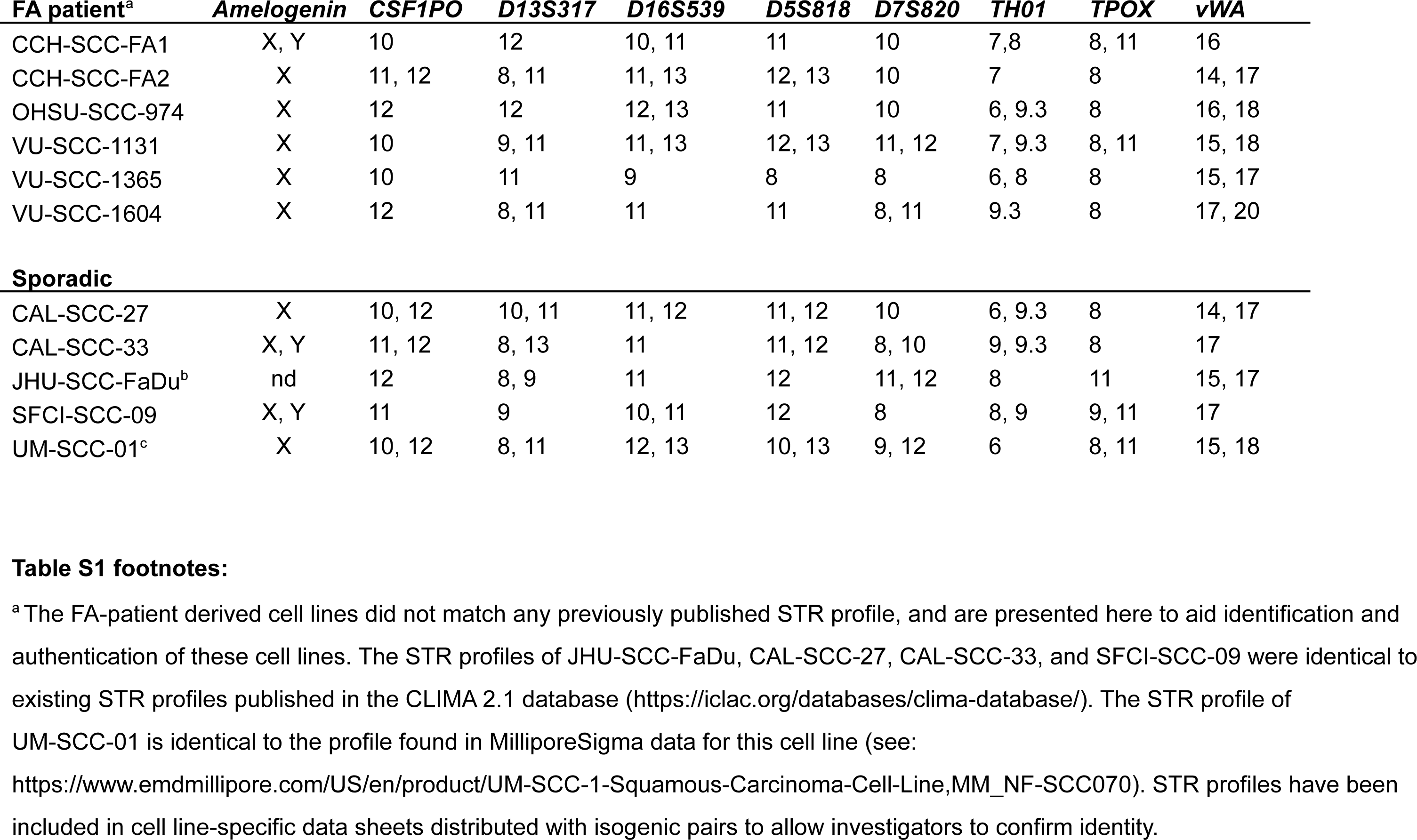
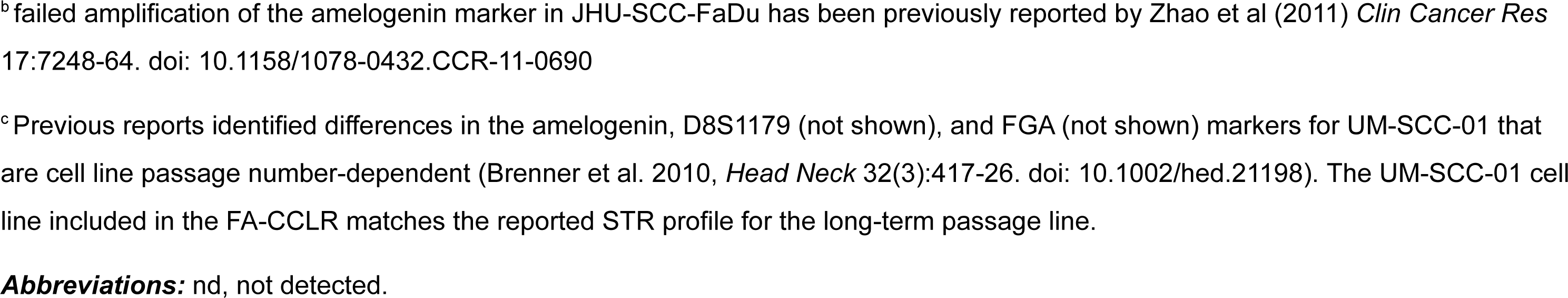
STR cell line authentication profiles for FA patient-derived and sporadic HNSCC cell lines.

| **FA patient<sup>a</sup>** | ***Amelogenin*** | ***CSF1PO*** | ***D13S317*** | ***D16S539*** | ***D5S818*** | ***D7S820*** | ***TH01*** | ***TPOX*** | ***vWA*** |
| --- | --- | --- | --- | --- | --- | --- | --- | --- | --- |
| CCH-SCC-FA1 | X, Y | 10 | 12 | 10, 11 | 11 | 10 | 7, 8 | 8, 11 | 16 |
| CCH-SCC-FA2 | X | 11, 12 | 8, 11 | 11, 13 | 12, 13 | 10 | 7 | 8 | 14, 17 |
| OHSU-SCC-974 | X | 12 | 12 | 12, 13 | 11 | 10 | 6, 9.3 | 8 | 16, 18 |
| VU-SCC-1131 | X | 10 | 9, 11 | 11, 13 | 12, 13 | 11, 12 | 7, 9.3 | 8, 11 | 15, 18 |
| VU-SCC-1365 | X | 10 | 11 | 9 | 8 | 8 | 6, 8 | 8 | 15, 17 |
| VU-SCC-1604 | X | 12 | 8, 11 | 11 | 11 | 8, 11 | 9.3 | 8 | 17, 20 |
| **Sporadic** |  |  |  |  |  |  |  |  |  |
| CAL-SCC-27 | X | 10, 12 | 10, 11 | 11, 12 | 11, 12 | 10 | 6, 9.3 | 8 | 14, 17 |
| CAL-SCC-33 | X, Y | 11, 12 | 8, 13 | 11 | 11, 12 | 8, 10 | 9, 9.3 | 8 | 17 |
| JHU-SCC-FaDu<sup>b</sup> | nd | 12 | 8, 9 | 11 | 12 | 11, 12 | 8 | 11 | 15, 17 |
| SFCI-SCC-09 | X, Y | 11 | 9 | 10, 11 | 12 | 8 | 8, 9 | 9, 11 | 17 |
| UM-SCC-01<sup>c</sup> | X | 10, 12 | 8, 11 | 12, 13 | 10, 13 | 9, 12 | 6 | 8, 11 | 15, 18 |

<sup>a</sup> The FA-patient derived cell lines did not match any previously published STR profile, and are presented here to aid identification and authentication of these cell lines. The STR profiles of JHU-SCC-FaDu, CAL-SCC-27, CAL-SCC-33, and SFCI-SCC-09 were identical to existing STR profiles published in the CLIMA 2.1 database (https://iclac.org/databases/clima-database/). The STR profile of UM-SCC-01 is identical to the profile found in MilliporeSigma data for this cell line (see: https://www.emdmillipore.com/US/en/product/UM-SCC-1-Squamous-Carcinoma-Cell-Line,MM_NF-SCC070). STR profiles have been included in cell line-specific data sheets distributed with isogenic pairs to allow investigators to confirm identity.
<sup>b</sup> failed amplification of the amelogenin marker in JHU-SCC-FaDu has been previously reported by Zhao et al (2011) *Clin Cancer Res* 17:7248-64. doi: 10.1158/1078-0432.CCR-11-0690
<sup>c</sup> Previous reports identified differences in the amelogenin, D8S1179 (not shown), and FGA (not shown) markers for UM-SCC-01 that are cell line passage number-dependent (Brenner et al. 2010, *Head Neck* 32(3):417-26. doi: 10.1002/hed.21198). The UM-SCC-01 cell line included in the FA-CCLR matches the reported STR profile for the long-term passage line.
**Abbreviations:** nd, not detected.

**Table S2.** *FANC*-isogenic cell line pair MMC IC_50_ values. The mitomycin-C IC_50_ nM concentration and 95% confidence interval (95% CI) values were determined for five FA patient-derived and three sporadic HNSCC cell line pairs where transgene complementation was performed by chromosomal safe harbor site transgene insertion (+Tg(SH) or by lentiviral transduction (+Tg(LV). Newly generated *FANCA* deletion sublines are indicated by ‘ko’ (knockout), followed by safe harbor site *FANCA* transgene insertion (+Tg). Fold differences as a function of genotype shown in the last column indicate either *FANC* transgene-complementation-enhanced MMC resistance (‘+’ values) or *FANCA* gene deletion-mediated MMC sensitization (‘-’ values) versus parental cell lines.

| Cell line | IC<sub>50</sub> (nM) | 95% CI (nM) | fold difference |
| --- | --- | --- | --- |
| CCH-SCC-FA1 | 4.65 | 2.93 - 5.96 |  |
| CCH-SCC-FA1+Tg(SH) | 16.17 | 13.43 - 19.18 | 3.48 |
| CCH-SCC-FA1+Tg(LV) | 47.12 | 42.29 - 52.52 | 10.13 |
| VU-SCC-1131 | 8.49 | 7.82 - 9.19 |  |
| VU-SCC-1131+Tg | 35.33 | 31.70 - 40.35 | 4.16 |
| VU-SCC-1365 | 3.05 | 0.89 - 4.97 |  |
| VU-SCC-1365+Tg | 22.74 | 21.29 - 24.32 | 7.47 |
| VU-SCC-1604 | 2.85 | 2.22 - 3.38 |  |
| VU-SCC-1604+Tg(SH) | 8.12 | 6.85 - 9.46 | 2.85 |
| VU-SCC-1604+Tg(LV) | 19.06 | 16.86 - 21.67 | 6.69 |
| OHSU-SCC-974 | 13.36 | 11.02 - 15.98 |  |
| OHSU-SCC-974+Tg(SH) | 39.77 | 35.14 - 44.92 | 2.98 |
| OHSU-SCC-947+Tg(LV) | 66.87 | 59.38 - 75.42 | 5.01 |
| JHU-SCC-FaDu | 32.38 | 28.40 - 36.92 |  |
| JHU-SCC-FaDu ko | 3.633 | 3.060 - 4.193 | -8.91 |
| JHU-SCC-FaDu ko+Tg | 33.54 | 31.65 - 35.53 | 0.96 |
| SFCI-SCC-09 | 27.37 | 20.72 - 40.60 |  |
| SFCI-SCC-09 ko | 0.62 | 0.10 - 1.42 | -44.37 |
| SFCI-SCC-09 ko+Tg | 19.22 | 17.81 - 20.77 | -1.42 |
| UM-SCC-01 | 36.80 | 31.87 - 42.45 |  |
| UM-SCC-01 ko | 2.64 | 2.05 - 3.26 | -13.92 |
| UM-SCC-01 ko+Tg | 36.15 | 31.57 - 41.44 | -1.02 |

**Table S3:**
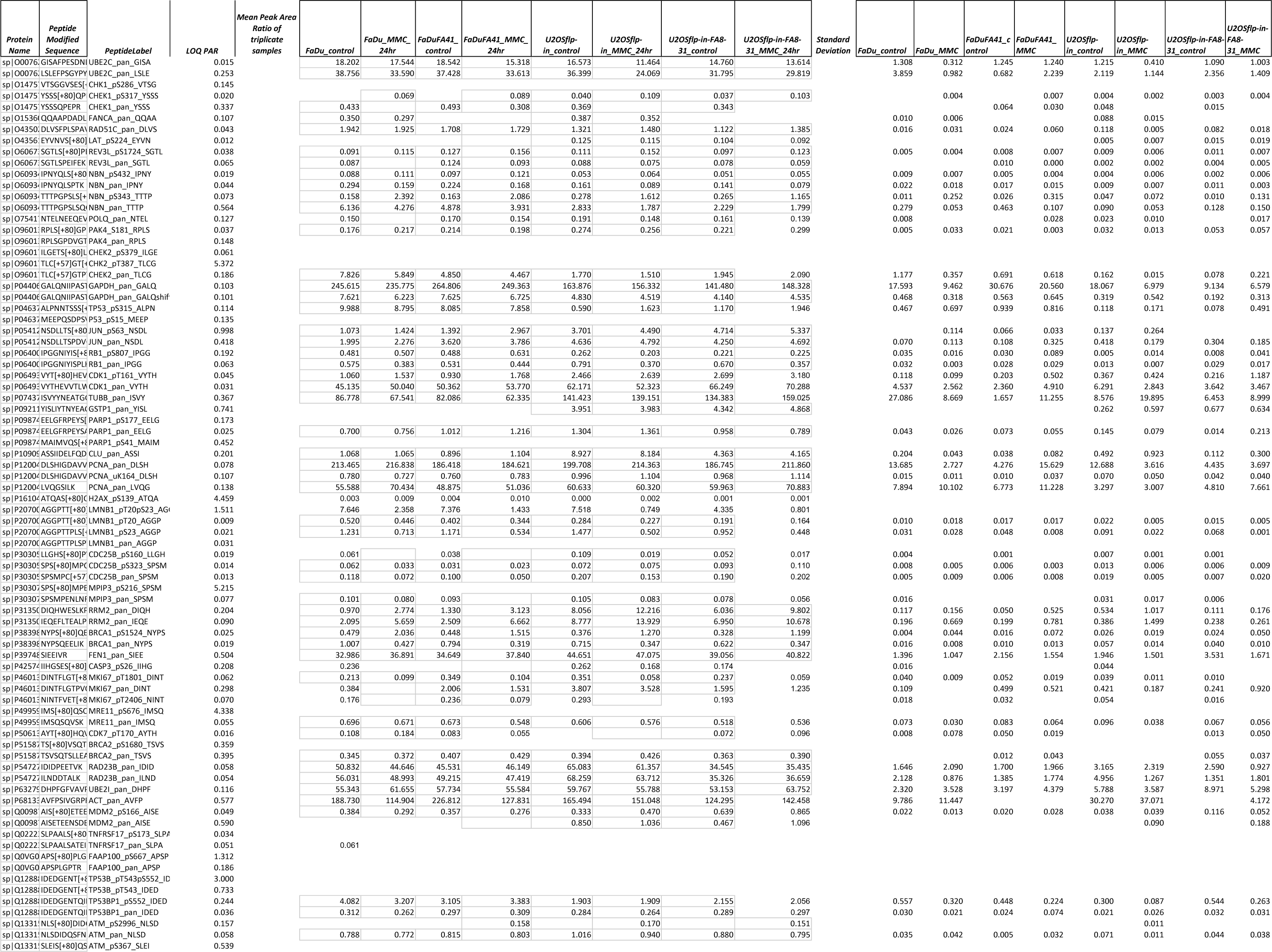
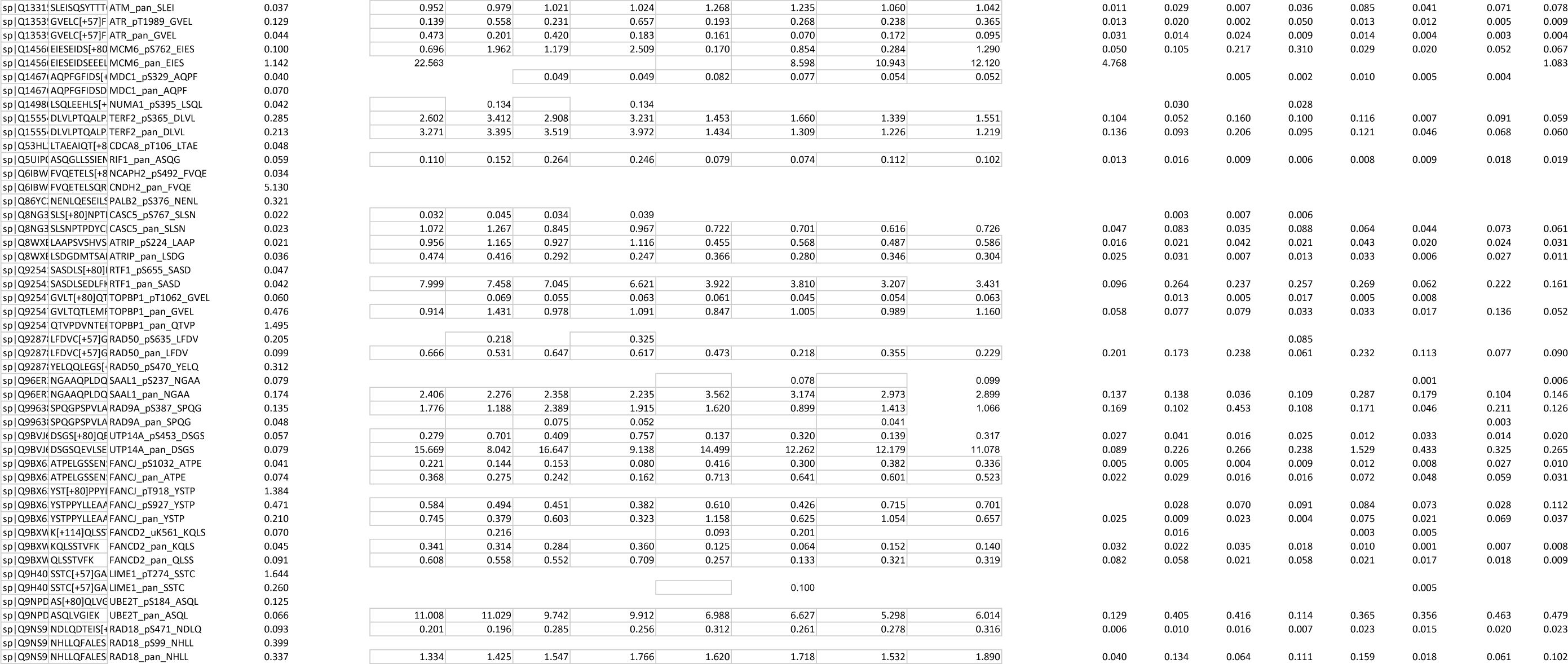
Mean peak area values for peptides detected by MRM-MS in MMC-treated isogenic pairs of control and *FANCA*-deficient JHU-SCC-FaDu and U-2 OS cell lines.

